# Finding recurrent RNA structural networks with fast maximal common subgraphs of edge-colored graphs

**DOI:** 10.1101/2020.02.02.930453

**Authors:** Antoine Soulé, Vladimir Reinharz, Roman Sarrazin-Gendron, Alain Denise, Jérôme Waldispühl

## Abstract

**Motivations:** RNA tertiary structure is crucial to its many non-coding molecular functions. RNA architecture is shaped by its secondary structure composed of stems, stacked canonical base pairs, enclosing loops. While stems are captured by free-energy models, loops composed of non-canonical base pairs are not. Nor are distant interactions linking together those secondary structure elements (SSEs). Databases of conserved 3D geometries (a.k.a. modules) not captured by energetic models are leveraged for structure prediction and design, but the computational complexity has limited their study to local elements, loops, and recently to those covering pairs of SSEs. Systematically capturing recurrent patterns on a large scale is a main challenge in the study of RNA structures.

**Results:** In this paper, we present an efficient algorithm to compute maximal isomorphisms in edge colored graphs. This framework is well suited to RNA structures and allows us to generalize previous approaches. In particular, we apply our techniques to find for the first time modules spanning more than 2 SSEs, while improving speed a hundredfold. We extract all recurrent base pair networks among all non-redundant RNA tertiary structures and identify a module connecting 36 different SSEs common to the 23S ribosome of *E. Coli* and *Thermus thermophilus*. We organize this information as a hierarchy of modules sharing similarities in their structure, which can serve as a basis for future research on the emergence of structural patterns.

**Availability:** http://csb.cs.mcgill.ca/carnaval2

## 1 Introduction

Functional RNA tertiary structures are stabilized by a collection of base pairs and base stackings often referred to as the secondary structure. The latter forms a planar structure made of stems of canonical base pairs (i.e. Watson-Crick and Wobble) connected by loops. Although these loops do not feature regular canonical base pairs patterns, they are often characterized by complex non-canonical base pair networks that create sophisticated 3D motifs used to shape the molecular structure. Furthermore, these loops occasionally interact with distant parts of the structure (i.e. other loops or stems) to form bridges stabilizing the global architecture of the RNA. The identification and characterization of these structural sub-units is therefore essential for a better understanding of the evolution of structured RNAs and the development of robust methods for predicting tertiary structures.

RNA modules are small and (usually) densely connected base pair patterns that can be observed in a variety of different molecules, sometimes in multiple locations. Fig. 1 displays an RNA secondary structure and, below, a module from the same structure to serve as an illustration. The conservation of RNA modules suggests an evolutionary pressure to preserve specific interaction patterns that constrains the possible set of sequences to the ones compatible with those interactions. As a consequence, identified RNA modules associate sequences to potential structures and so help to draw information about base pairs out of RNA sequences. This information can then be used to infer the 3D structure of the whole molecule [11, 15, 16, 14, 20, 13, 21].

**Figure 1:**
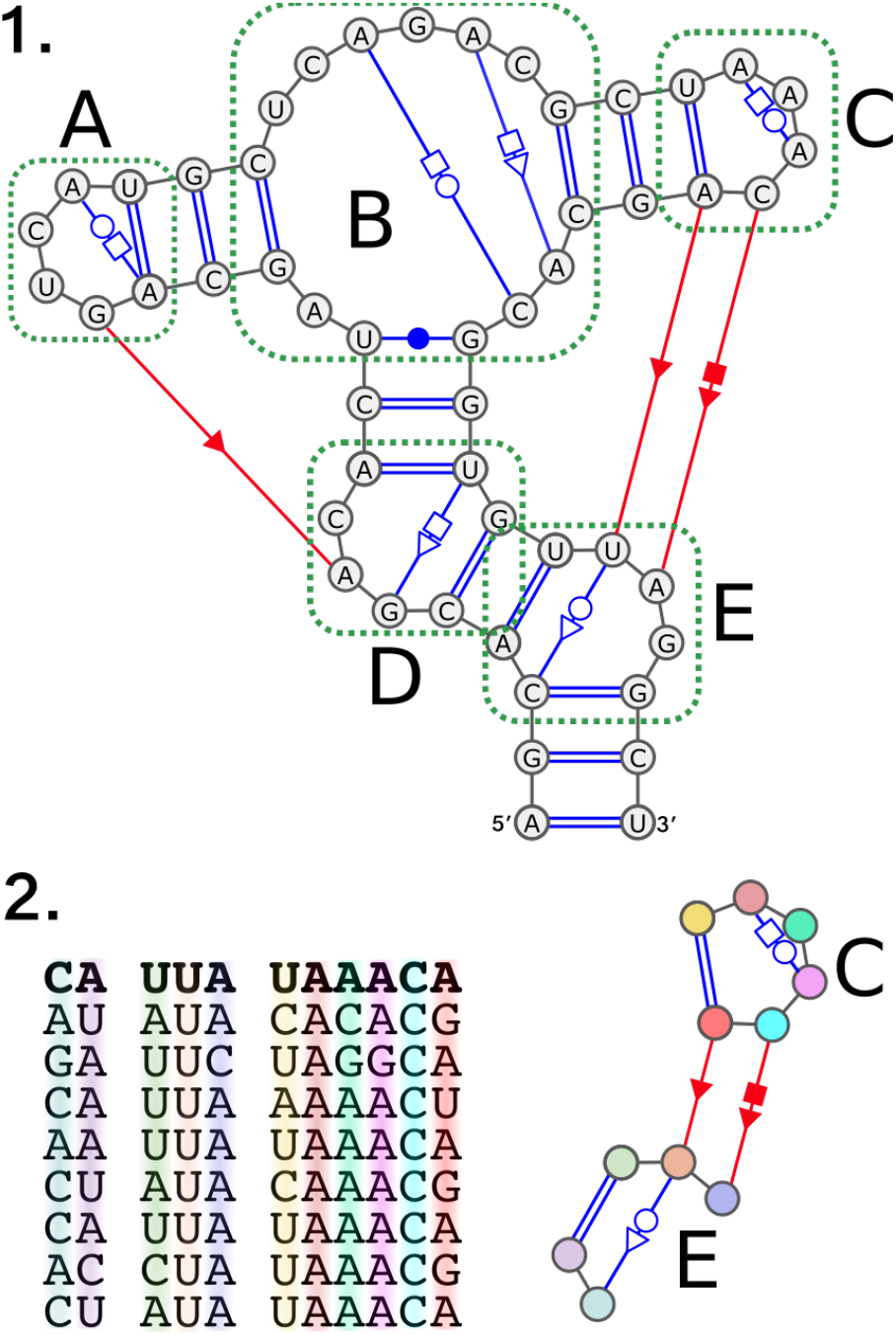
Secondary structure and module. In (1) we show an RNA and its secondary structure with non-canonical interactions. Base pair interactions in blue are local (both nucleotides involved are in the same or in adjacent SSEs) while the ones in red are long range interactions (between two distant SSEs). The canonical base pair interactions are represented with double lines. We highlighted the loops in the structure with green dotted lines. Loops **A** and **C** are hairpins, loops **D** and **E** are interior loops, and loop **B** is a multi-loop. In (2) we show an instance of a module found in the RNA secondary structure in (1). On the right is the base pair pattern that characterizes this module and on the left is the sequence profile of this module (i.e. the nucleotide sequences of the corresponding parts of RNAs this module has been observed in). The first sequence in the profile, for instance, corresponds to the RNA displayed in (1).

Other applications require a well defined and rigorous description of modules. In synthetic biology, the availability of databases of autonomous structural modules is key for designing new molecules [29]. The assembly of RNA binding sites may also require bringing together distant modules within the secondary structure [22]. A comprehensive and indexed catalog of sub-structures would greatly facilitate studies of these sites.

Some RNA modules have received a specific attention such as *GNRA loops, Kink-turns, G-bulges*, and the various types of *A-minors*. Moreover, several works have been presented, proposing computational methods to detect RNA modules in tertiary structures using either geometry or graph-based approaches [1, 5, 6, 7, 8, 9, 19, 24, 26, 28, 30, 4, 2]. However, the purpose of the majority of those methods is to search for known modules in new structures. A couple of methods has been proposed that search local modules without any prior knowledge of their geometry or topology [5, 9]. In addition to those methods, databases of RNA modules found in experimentally determined RNA tertiary structures have been proposed such as RNA 3D Motif Atlas [20] and RNA Bricks [3].

We are interested in the whole landscape of RNA modules (known or not) rather than any RNA module in particular which distinguishes us from most of the works previously mentioned. Furthermore, we aim at extracting recurrent patterns in the secondary structure rather than in the sequence or in the tertiary structure. Indeed, those patterns capture topological information that implies a similar tertiary structure and a consensus RNA sequence can be derived from it. As such they constitute interesting RNA modules candidates. Our goal is to automatically capture this topological information.

To our knowledge, the only published method similar in those aspects is `CaRNAval` [23], one of our previous work. In `CaRNAval`, we presented an algorithm to find all identical *interaction networks* between two RNAs [23], which capture the topological information of interaction modules (i.e. RNA modules over two, non-adjacent, secondary structure elements or SSEs) but not the sequences. We made the results of `CaRNAval` available as an extensive organized catalogue of the *Recurrent Interaction Networks* (RINs) computed on all the non-redundant structures available in RNA3DHub [18]. The method developed for `CaRNAval` is automated and does not use any prior knowledge of neither the topology nor the geometry of the structures it detects.

Compared to `CaRNAval`, the work presented in this paper takes several steps toward capturing the whole landscape of RNA modules. Indeed, by approaching RNA secondary structures as graphs equipped with a *proper edge coloring*, we designed several graph matching algorithms and used them as the core of a modular automated pipeline. Leveraging the *proper edge coloring* of a structure graph allows to improve execution time a hundredfold compared to `CaRNAval`. Moreover, and this is the main novelty of this method, there are no built-in constraints on the structures it can capture (albeit it accepts such constraints as an optional input). This flexibility joined with the improved performances allow to mine for any kind of RNA module candidates.

Typically, our method can capture structures spanning an arbitrary large number of SSEs when all other approaches are only considering similarities between a loop and `CaRNAval` only extended this analysis to pairs of loops connected together. We can thus compute similarities between arbitrarily large RNAs (the largest module candidate we observe spans 293 nodes with 610 edges). Moreover, we show that the new structures found by removing this restriction complement the landscape of modules presented in `CaRNAval` and so are other new structures obtained by broadening the search space further. As a consequence, our results underline the universality and fundamental nature of these recurrent architectures.

## 2 Method

From a set of *mmCIF* files describing 3D structures of RNA chains, we first annotate the interactions with `FR3D`. The method presented analyze these annotations in four steps.

1. We first build for each chain a directed edge-labelled graph such that the edges represent the phosphodiester bonds as well as the canonical and non-canonical interactions. The labels on the edges correspond to the interaction types plus the indication of the interaction being either local (inside a single SSE) or long-range (between two SSEs)
2. For each pair of RNA graphs, we extract all the Maximal Common Subgraphs such that edges are matched to edges with the same labels
3. Each Maximal Common Subgraph is then processed to obtain the Recurrent Structural Elements (constrained common subgraphs) it contains
4. Finally we gather the Recurrent Structural Elements found together into a non-redundant collection and create a network of direct inclusions.

### 2.1 RNA 2D Structure Graphs

We rely on RNA 2D structure graphs to represent the structures of RNA chains. RNA 2D structure graphs are directed edge-labelled graphs. Each node represents a nucleotide, each edge represents an interaction (base pair or backbone). Edges are labelled according to the annotation of the interaction they correspond to. Annotations of base pair interactions follow the Leontis-Westhof geometric classification [12]. They are any combination of the orientation cis (c) (resp. trans (t)) with the name of the side which interacts for each of the two nucleotides: Watson-Crick (W, represented with ● in cis orientation or ○ in trans), Hoogsteen (H, ■ in cis □ in trans) or Sugar-Edge (S, ▶ in cis ▷ in trans). Thus, each base pair is annotated by a string from the set: {c,t}×{W,S,H}^2^ or by combining the corresponding symbols. Note that canonical cWW interactions constitute an exception and are represented with a double line instead of “● ●”. Moreover, each basepairs interaction can also be annotated as either *local* or *long range*, depending on the secondary structure elements the nucleotides involved are found in (our method to generate the secondary structure is described in section 3.1). The backbone is represented with directed edges, labelled *b*53.

As a consequence, an annotation (and thus an edge label) is composed of three characters *xYZ* ∈ [*c* | *t*][*W* | *S* | *H*]^2^ plus a parameter *C* ∈ [local | long-range]. Interactions are either symmetric (*xYY*) or not symmetric (*xYZ*). Each non symmetric interaction between nucleobases *xYZ* is complemented by an interaction *xZY* between the same nucleobases and the same value of *C* but in the opposite direction. We introduce an abstract type/label *b*35 to complement the *b*53 label. We can thus define a bijection *ι* as follow:

- *ι*(*xYZ, C*) = *xZY, C*
- *ι*(*xYY, C*) = *xYY, C*
- *ι*(*b*53, local) = *b*35, local
- *ι*(*b*35, local) = *b*53, local

An interaction of type *t* between nucleotides *a,b* (represented by nodes *υ_a_,υ_b_*), is represented by two directed edges {*υ_a_,υ_b_*} and {*υ_b_,υ_a_*} whose respective labels are *t* and *ι*(*t*). This property is important as a requirement of the algorithms we designed (cf. section 1.5.1 of the supplementary material).

We represent each RNA chain in the dataset as a RNA 2D structure graph, the annotations of the RNA base pair interactions corresponding to the labels of the edges of the graph (cf. Fig. 2).

**Figure 2:**
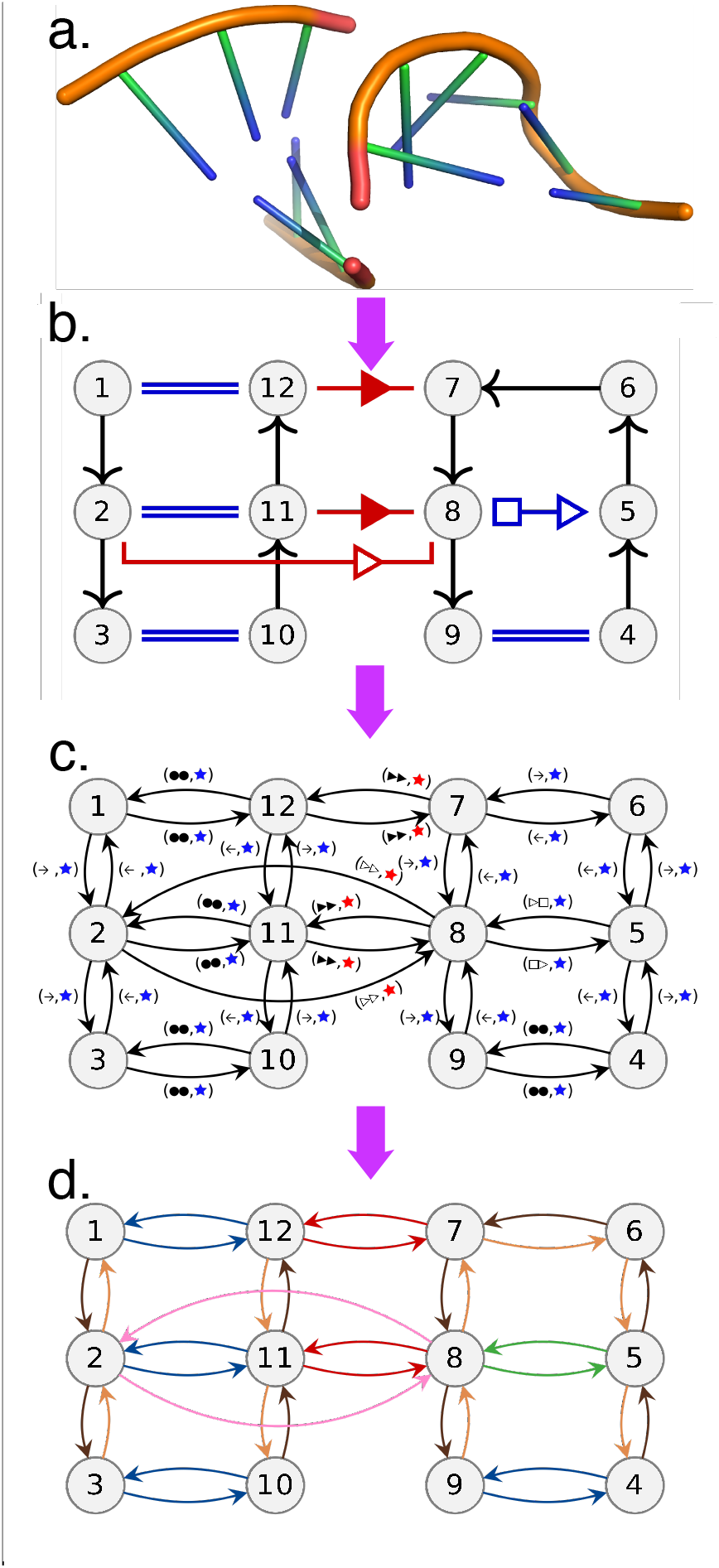
From 3D structure to directed edge-labelled graph. In this figure we illustrate the transition from the 3D structure (a) to RNA 2D structure graph (b) and finally directed edge-labelled graph (c) with a simple RNA structure. Each edge label of the directed edge-labelled graph is a pair which first element represents the type of interaction (using the same symbols as in the RNA 2D structure graph) while the second denotes the local (blue) vs. long-range (red) property of the interaction (using the same colors as in the RNA 2D structure graph). Moreover, the set of edge labels forms a directed *proper edge-coloring*, as illustrated with the last panel (d).

### 2.2 Graph Matching & Proper Edge-Coloring

As we transpose RNA structures into edge-labelled graphs, finding common substructures in the RNA structures comes down to finding common subgraphs in the RNA 2D structure graphs.

Problems that consist in matching graphs or parts of graphs are called *Graph Matching* problems. We are especially interested in finding common subgraphs, an NP-hard problem in general. However, RNA 2D structure graphs inherit some of the constraints of the RNA structures they represent, constraints that translate into a graph property useful for graph matching.

The chemical constraints of nucleotides interactions are such that each edge of a nucleotide should be involved in at most one interaction. This translates in terms of graphs as follows: for all RNA 2D structure graphs *G* = {*V, E*} and for all a node *υ* ∈ *V*, there are no two edges *e*_1_, *e*_2_ ∈ *E* that originate from *υ* with the same label. To put it differently, the set of labels on the edges of any RNA 2D structure graphs naturally forms a *Proper Edge-Coloring* (PEC). We designed three graph matching algorithms designed to take advantage of the proper edge-coloring the RNA 2D structure graphs come equipped with.

### 2.3 Exceptions

We observed a few nucleotides annotated with two interactions involving the same Leontis-Westhof edges in some RNA structures (0.02% of the nucleotides of our reference dataset cf. section 3.1). Those interactions could either be annotation errors or biologically relevant. Given the rarity of those exceptions, we chose to duplicate the graphs concerned into different proper edge-colored versions, each covering a different interpretation. Details about the duplication procedure and the different versions are provided in section 2.1 of the supplementary material.

### 2.4 Graph Matching Algorithms

In this section we briefly introduce our 3 algorithms, the 3 problems they solve and how we take advantage of the PEC. Extensive and formal descriptions are provided in the supplementary material (sections 1.2, 1.3 and 1.4).

#### 2.4.1 Definitions & Notations

Two graphs *G* = {*V_G_, E_G_*} and *H* = {*V_H_,E_H_*} are isomorphic *iff* there is a bijection *b* from *V_G_* to *V_H_* that respects the edges and their labels. A graph *G* = {*V_G_, E_G_* } is a *subgraph* of graph *H* = {*V_H_, E_H_* } *iff* there exists at least one injection *i* from *V_G_* to *V_H_* that respects the edges and their labels.

Given two graphs *G,H*, a graph *S* = (*V_S_,E_S_*) is a *common subgraph* of *G* and *H* if it is a subgraph of *G* and a subgraph of *H*. A common subgraph *S* of *G* and *H* is *maximal iff* for all *S*′ subgraph of *G* and *H, S* ⊂ *S*′ ⇒ *S* = *S*′. All three algorithms take two properly edge-colored graphs *G* = {*V_G_, E_G_*} and *H* = {*V_H_, E_H_*} as an input. For any color *c*, the sets of c-colored edges are denoted *E_Gc_* and *E_Hc_*.

#### 2.4.2 Using the PEC when extending a matching

The three algorithms presented in this paper revolve around exploiting the constraint added by having to respect the PEC when matching two graphs to greatly reduce the search space. All three algorithms reliy on the same core strategy. Matching the two graphs is done by starting with a minimal match and then extending it through the neighbors of the already matched nodes. This strategy is common in graph matching and usually requires to test all permutations between the two sets of neighbours. However, the constraint of respecting the PEC only leaves at most a single valid affectation of the neighbours, as illustrated in figure 3. As a consequence, the complexity of the extension process is linear in the number of nodes (since the number of colors is fixed, cf. section 1.2.3 of the the supplementary material).

**Figure 3:**
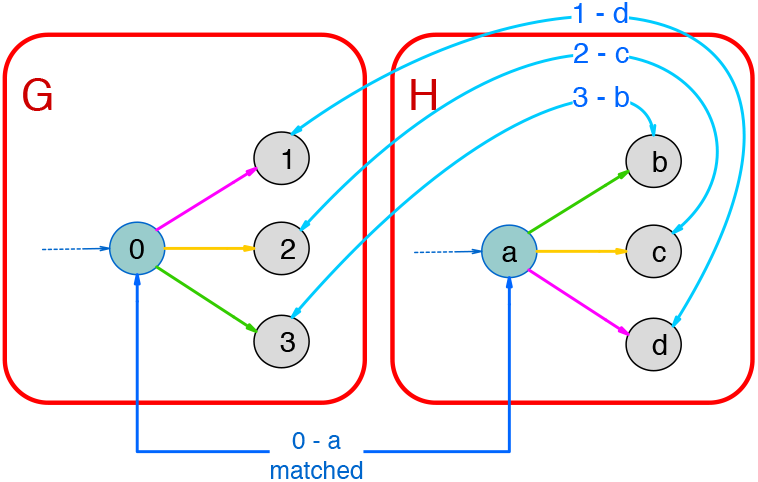
Impact of proper edge-coloring on graph-matching. This figure displays a piece of two graphs (G on the right and H on the left) in which the nodes 0 and *a* are already matched together. The next step is to match their neighbours. In the generic case, all permutations have to be tested. On the contrary, in the example displayed, the colors of the edges limit the options to consider to a single one.

#### 2.4.3 Graph Isomorphism Algorithm

The *Graph Isomorphism* problem consists in determining if two properly edge-colored graphs *G* and *H* are isomorphic. Our Graph Isomorphism Algorithm determines the color *c* that minimizes the product |*E_G,c_*| × |*E_H,c_*|. Then, for all pairs of edges ({*g*_1_, *g*_2_}, {*h*_1_, *h*_2_}) ∈ *E_G,c_* × *E_H,c_*, the algorithm launches an extension with the matching ((*g*_1_, *h*_1_), (*g*_2_, *h*_2_)) as starting point. The two graphs are isomorphic *iff* it exists a matching that can be extended into a bijection of *V_G_* and *V_H_* that respects the edges and their coloring. As we mentioned previously, the extension process is in 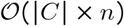 (assuming *n* = |*V_G_*| = |*V_H_*|, if not, *G* and *H* are trivially not isomorphic) and the number of starting point is capped by 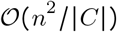 resulting in a 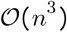 complexity for the algorithm (cf. section 1.2.3 of the the supplementary material).

#### 2.4.4 Subgraph Isomorphism Algorithm

The *Subgraph Isomorphism* problem consists in, given two properly edge-colored graphs *G* and *H*, determining if *G* is a subgraph of *H*. Our Subraph Isomorphism Algorithm is derived from our Graph Isomorphism Algorithm, the difference between the two being that *G* is a subgraph of *H iff* it exists a matching that can be extended into an injection of *V_G_* in *V_H_* that respects the edges and their coloring. The complexity is the same as the Graph Isomorphism Algorithm: 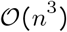 with *n* = *min*(|*V_G_*|, |*V_H_*) (cf. section 1.3.3 of the the supplementary material).

#### 2.4.5 All Maximal Common Subgraphs Algorithm

The *All Maximal Common Subgraphs* problem consists in finding all maximal common subgraphs between two properly edge-colored graphs *G* and *H* (note that this differs slightly from the *maximal common subgraph* problem which usually consists in just finding the largest common subgraph). This algorithm relies on the same extension strategy than the two previous ones. However, unlike the two previous problems, encountering a discrepancy during the extension does not imply that this extension should be abandoned (as illustrated in Fig. 4). Instead, it suggests the existence of an alternative way of matching the graphs by considering the nodes in a different order than in the current extension. As we are looking for all maximal common subgraphs, this alternative has to be explored as well. As a consequence, we designed an unconventional backtracking mechanism. For any new discrepancy encountered, we launch a new extension with a list of constraints (similar to instructions) designed to force this new extension to explore the alternative suggested by the discrepancy. Such an extension can also encounter new discrepancies and so on and so forth. Figure 5 illustrates this process and a complete description of this mechanism (with additional illustrations) is provided in section 1.4.2 of the supplementary material as well as a formal proof of its correctness in section 1.4.3.

**Figure 4:**
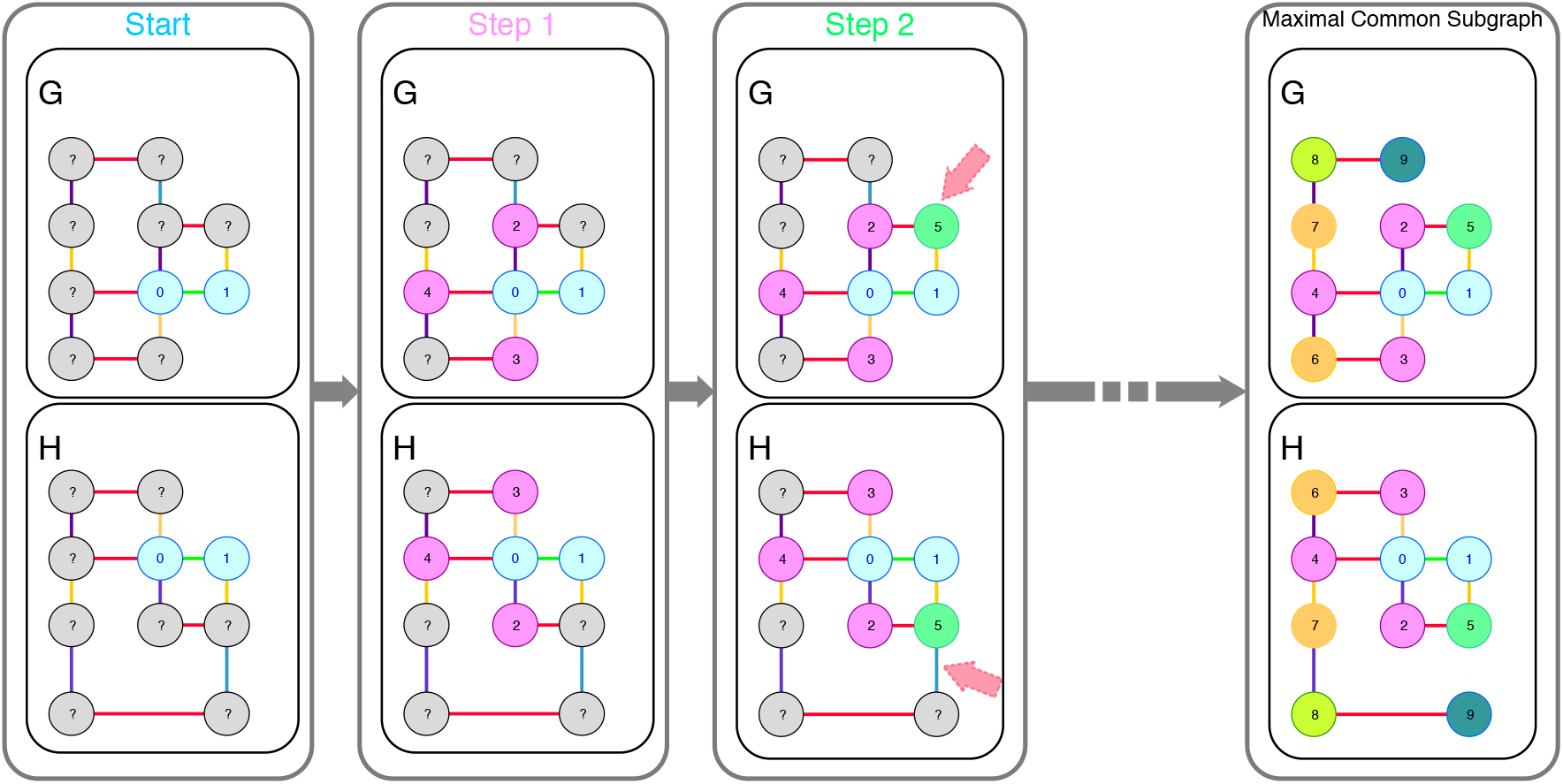
Illustration of the extension process. This figure illustrates the extension process from a “starting point” (here ((*g*_0_, *h*_0_), (*g*_0_, *h*_0_)), in blue). We first consider the neighbors of *g*_0_ and *h*_0_ (in purple). Thanks to the PEC, there is only one way to match them. We then consider the neighbors of *g*_1_ and *h*_1_ (in green). We match *g*_5_ and *h*_5_ but discover that their neighborhoods are not compatible. At this point the behaviours of the three algorithms differ. This discovery implies that the matching cannot be extended to cover all of *G* so the *Graph Isomorphism* and *Subgraph Isomorphism* will abandon it and pass on to another “starting point”. The *All Maximal Common Subgraphs* on the contrary will take note of this discrepancy and keep extending the matching nevertheless. This extension will output a maximal common subgraph of *G* and *H* and a new branch will be created to explore the alternative solution suggested by the discrepancy found.

**Figure 5:**
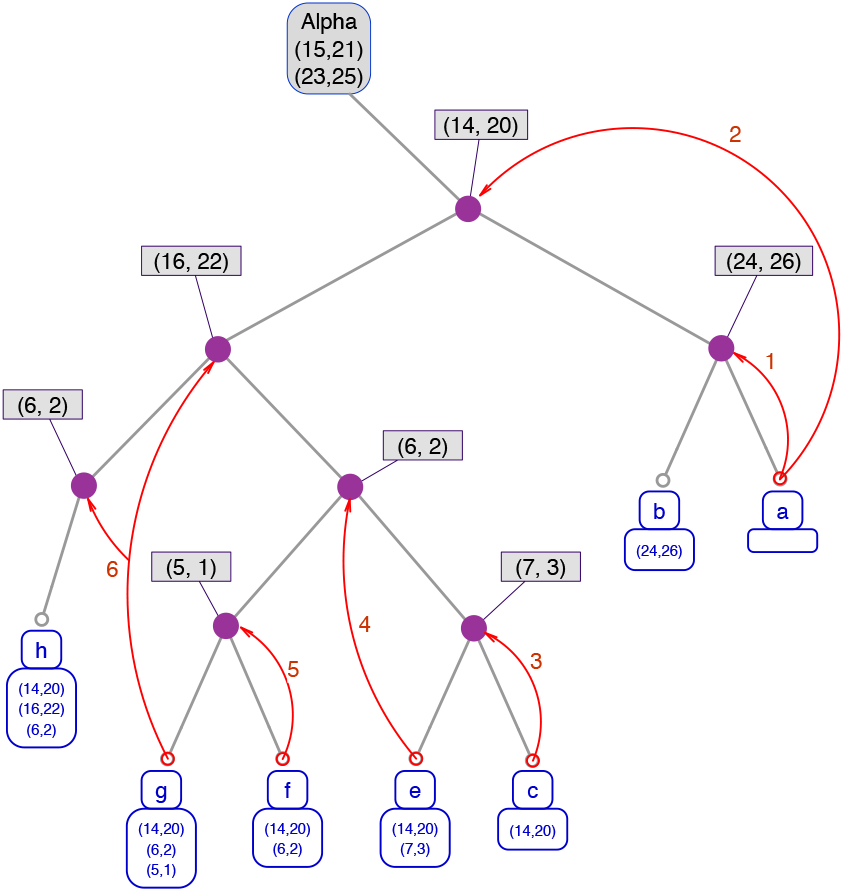
Exploration tree with backtracking. This figure displays the exploration tree representing *a posteriori* the relation between the different branches created. In this tree, the root is a starting point (i.e. the nodes that are already matched at the start of an exploration) and each leaf is a different maximal common subgraph. Each path from the root to a leaf describes an exploration. For instance, the node (14,20) of the exploration tree corresponds to the action of matching the node 14 from G to the node 20 of H. All the leafs in the right subtree have matched 14 to 20 and all the ones in the left subtree have not. Note that only the nodes with a left child are represented, all other nodes have been collapsed since they bear no information about the exploration process. The first exploration always produces the right most maximal common subgraph. In this exemple, the first exploration encountered two conflicts and the algorithm thus produced two new branches which respectively were instructed not to add (24,26) and not to add (14,20). The first of the two produced another maximal common subgraph without any trouble but the second encountered another conflict and so on and so forth.

### 2.5 From common subgraphs back to RNA structures

By transposing the RNA structures to graphs and using our algorithms, we are thus able to obtain the set of *All Maximal Common Subgraphs* contained in any given dataset. However, the size of this set grows exponentially with the size of the dataset, quickly making it humanly unmanageable. As a consequence, we designed a restriction system to define more human-sized subsets of structural elements and designed our method to extract and organize such subsets specifically rather than the whole set *All Maximal Common Subgraphs*. Those subsets of structural elements are to be defined by users through rules or restrictions, according to the types of structures they want to study.

One of the strong points of our methods is its ability to easily switch from a subset to another since the restriction system is independent from the graph matching part. This opens the opportunity to conduct studies on several related subsets to draw comparisons, as illustrated in section 3. Since we will be working on different subsets simultaneously, let us formalize what those subsets are or can be.

#### 2.5.1 Recurrent Interaction Network (RIN)

We call *Recurrent Interaction Network* (RIN) any recurrent subgraph of RNA 2D Structure Graphs (i.e. observed in at least two RNAs of the dataset). A RIN is formally defined as a pair 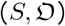 with:

- *S* = {*V_S_, E_S_*} a connected graph with the properties of a RNA 2D structure graph
- 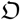 a collection of *occurrences*. An *occurrence* records an observation of *S* in the dataset. We represent an *occurrence* as a pair (*G, i*) with *G* = {*V_G_, E_G_*} a RNA 2D structure graph and *i* an injection from *V_S_* to *V_G_* that respects the edge labels.
- 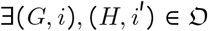 s.t. *G* ≠ *H* (i.e. it should be *recurrent*)

This minimal set of properties defines the *RIN*\* class^1^ which can be seen as the mother-class from which all other classes are derived by adding additional restrictions.

To illustrate this let us consider a set of additional rules/restrictions R, designed to invalidate some structural elements we are not interested in. *R* thus defines *RIN^R^* which is a subclass of *RIN*\*. For our method to extract *RIN^R^* from a dataset, R is to be translated into a filtering function *f_R_*: *G* → *C_RIN^R^_* with *G* a graph that shares the same properties as an RNA 2D structure graph and *C_RIN^R^_* the collection of 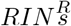 in *G* that respects the rules in *R* (the properties defining *RIN*\* are “built-in”). To put simply, the role of *f_R_*: *G* → *C_RIN^R^_* in the pipeline is to extract the 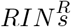 from the maximal common subgraphs.

Additionally, we offer the possibility of providing a second filtering function 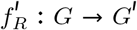 that takes as input an RNA 2D structures graphs *G* in the dataset and outputs another graph *G*′, which is a subgraph of *G* without the edges and nodes in *G* that already infringe a rule of *R* (and thus cannot possibly be part of any valid *RIN^R^*). 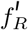 is optional as it only improves performances by reducing the search space, albeit greatly in most cases.

We will be using *RIN^R^* in the following sections to denote an arbitrary class of RINs currently being extracted.

#### 2.5.2 Extraction of *RIN^R^*

For every pair of RNA 2D Structure Graphs in the dataset (after the application of 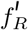, if provided), we use our algorithm solving the *maximal common subgraphs* problem to extract the set of all maximal common subgraphs between the two graphs (as illustrated in Fig. 6). The filtering function *f_R_* (derived from the rules in *R* that defines the class *RIN^R^* currently being extracted) is applied to each maximal common subgraph found. The sets of RINs obtained are gathered and clustered using our *graph isomorphism* algorithm. This process involves non trivial but incidental mechanisms which we describe in section 2.2 of the supplementary material.

**Figure 6:**
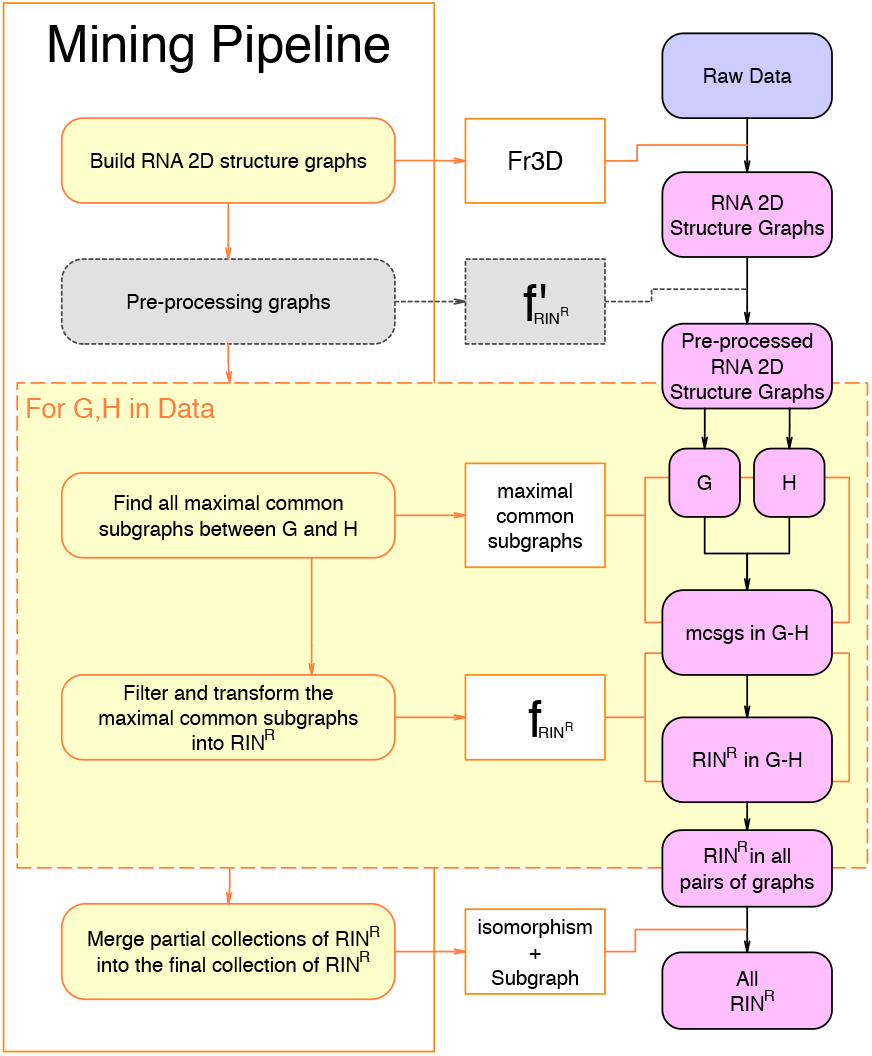
Simplified display of the full pipeline. The RNA 2D structure graphs given as input are pre-processed for the sake of optimization. Each pair of graphs in the pre-processed data is then given to the maximal common subgraphs algorithm as input and the output is post-processed into partial sets of 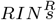. All partial sets of 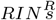 are finally merged into the complete set of 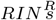 which is the output of the whole pipeline.

Note that our implementation relies on parallelization to improve the performances by distributing the pairs of graphs to process (cf. section 2.3 of the the supplementary material).

#### 2.5.3 Network of *RIN^R^*

RINs of a given class are often related (i.e. the canonical graph of one may be a subgraph of the canonical graphs of one or several others RINs). In order to display the internal structure of a class of RINs, we organize it into a network *N* = {*V, E*}. A node in *V* represents a RIN. An edge *e* = {*r*_1_, *r*_2_} from RIN 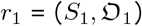 to RIN 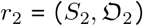, is in *E* iff *S*_1_ is a subgraph of *S*_2_. If the network is to be displayed, we then remove any edge *e* = {*r*_1_, *r*_3_} ∈ *E* if *e* = {*r*_1_, *r*_2_} ∈ *E* and *e* = {*r*_2_, *r*_3_} ∈ *E* to avoid overloading the display as the edges removed were equivalent to paths in the new version of the network. We rely on our *subgraph isomorphism* algorithm to build those networks efficiently.

## 3 Applications & Results

In this section, we present three applications of our method to three different yet related classes of RINs and the corresponding results.

### 3.1 Dataset

All three applications use the same dataset of RNA structures: the non-redundant RNA database maintained on RNA3DHub [18] on Sept. 9^th^ 2016, version 2.92. It contains 845 all-atom molecular complexes with a resolution of at worse 3Å. From these complexes, we retrieved all RNA chains also marked as non-redundant by RNA3DHub. The basepairs were annotated for each chain using `FR3D`. Because `FR3D` cannot analyse modified nucleotides or those with missing atoms, our present method does not include them either. If several models exist for a same chain, only the first one was considered.

To distinguish between local and long-range interactions, we define a secondary structure from the ensemble of canonical CWW interactions. This task can be ambiguous for pseudoknotted and large structures. We used the K2N algorithm [27] from the PyCogent library [10]. A case that can not be treated by K2N is when a nucleotide is annotatedas having two CWW interactions. Since this is rare, we decided to keep the interaction belonging to the largest stack.

### 3.2 Three different yet related classes of RINs

In this section we study three classes of RINs which are successive generalizations obtained by incrementally relaxing rules. As a consequence, we will first introduced those rules before introducing the different RIN classes we will be working on.

For any 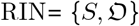, where *S* is a *canonical graph* representing the interactions network while 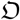 is the collection of occurrences:

*x* - each node in the canonical graph *S* belongs to a cycle in the undirected graph induced by *S*. (The undirected graph induced by *S* is obtained by replacing every directed edge by an undirected edge and merging those between the same nodes.)
*y* - if two nodes, *a* and *b* in *S*, form a local canonical base pair, there exists a node *c* in *S* such that *c* is a neighbor to *a* or *b*, and *c* is involved in a long-range or non-canonical interaction. In other words we do not extend stacks which nucleotides are involved in canonical base pairs only.
*z* - each node in *S* is involved in a canonical or a non-canonical interaction (*i.e*. no nodes with only backbone interactions)
*b* - *S* contains at least 2 long-range interactions, i.e. 4 edges labeled as long-range since each interaction is described with two directed edges.
*c* - the nucleotides corresponding to the nodes in *S* are captured by exactly 2 SSEs.

Rule *x* aims at enforcing the cohesiveness of the interaction network by preventing danglings that would create variations of little interest. Rule *y* aims at excluding pure stacks of canonical base pairs (i.e. at least two consecutive cWW with no other interaction) which form the core of the structure and are either embedded in the secondary structure with little geometric variation or result from the folding of the tertiary structure (co-axial stacking between helices, loop-loop interactions or pseudo-knots) with often a larger geometric variation. Rules *z* aims at excluding non interacting nucleotides that do not have geometric constraints as interaction networks are intended to capture a representation of the geometry. We will discuss the two last rules in parallel of the description of the classes.

We will denote the different RIN classes by concatenating the symbols of the rules that defines them (for instance *RIN^xyz^* is the class defined by the first three rules). This naming system has the advantage of making the name of a class an exact description of its definition. However, since the rules *x,y*, and *z* will be common to all classes, **we will replace xyz with a** in classes names. Please refer to table 1 for a summary of the different classes, their names and the rules they enforces. We also provide examples of structures in table 2 to illustrate how the successive relaxations of rules allow additional structures to be captured.

**Table 1:** Summary of the relation between the rules and the three RIN classes.

| Rules↓ | Classes→ | $RIN^{abc}$ | $RIN^{ab}$ | $RIN^a$ |
| --- | --- | --- | --- | --- |
| $a$ | Each node is in a cycle | ✓ | ✓ | ✓ |
|  | Stems of canonical base pairs are not extended | ✓ | ✓ | ✓ |
|  | Each node forms at least one base pair | ✓ | ✓ | ✓ |
| $b$ | At least two long range interactions | ✓ | ✓ | - |
| $c$ | The entire RIN must be over exactly two SSEs | ✓ | - | - |

**Table 2:**
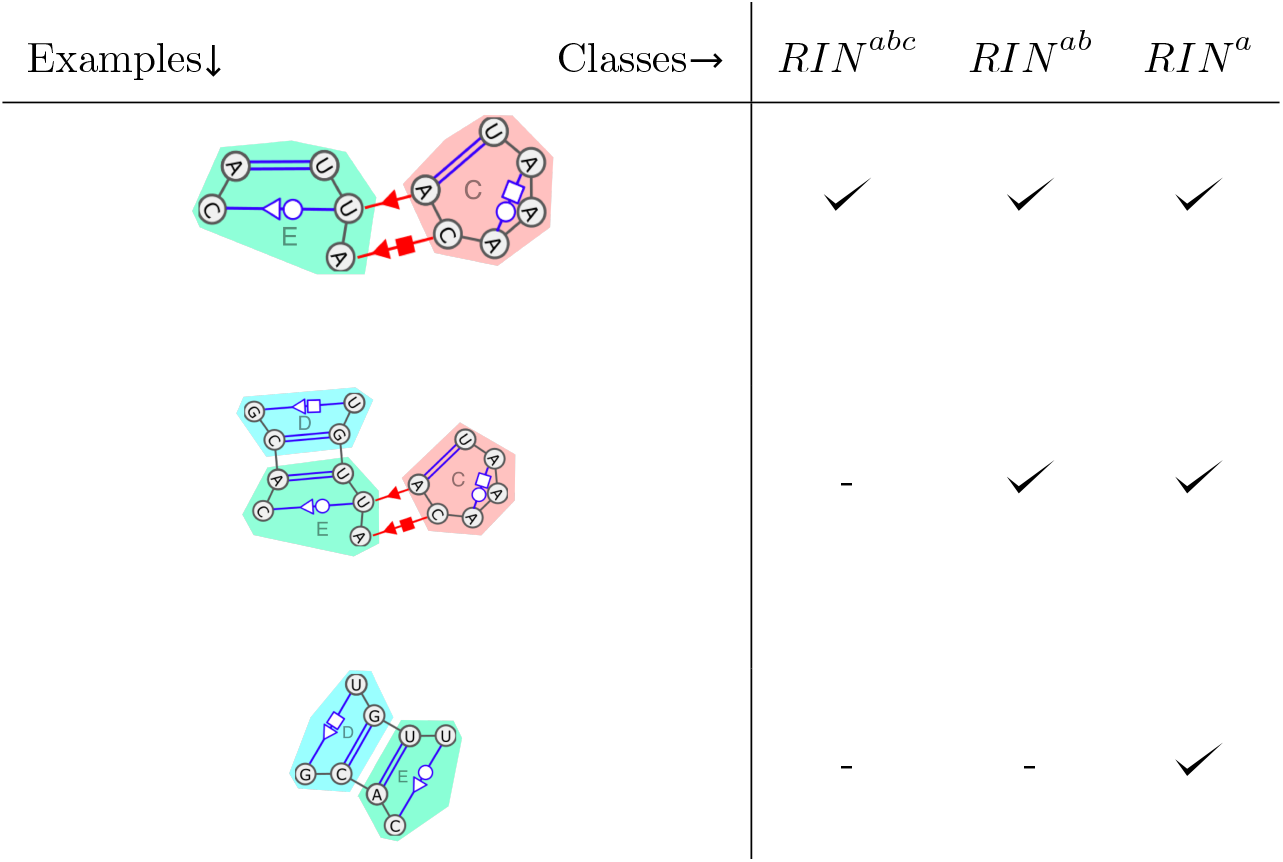
Examples of structures to illustrate the three RIN classes. Those three graphs are subgraphs of Fig. 1 with the same SSEs annotations (SSEs D,C and E figured with colored areas). The first graph is valid for all three classes. The second is over 3 SSEs and so cannot be a valid *RIN^abc^*. The third does not contain long-range interactions and thus is only valid for class *RIN^a^*.

| Examples↓ | Classes→ | $RIN^{abc}$ | $RIN^{ab}$ | $RIN^a$ |
| --- | --- | --- | --- | --- |
|  |  | ✓ | ✓ | ✓ |
|  |  | - | ✓ | ✓ |
|  |  | - | - | ✓ |

We inherit those five rules from the `CaRNAval` project [23]. The `CaRNAval` project aimed at extracting RNA structural motifs containing non-canonical base pairs, 2 or more long range interactions and involving exactly 2 SSEs. The set of structures extracted in `CaRNAval` corresponds in our system to the *RIN^abc^* class. We will use the *RIN^abc^* class as the reference and validation of our methods.

The second class we are working on is a generalization of *RIN^abc^* obtained by relaxing rule c and is thus called the *RIN^ab^* class. Rule *c* (having exactly 2 SSEs) was partially the consequence of the limits of the algorithm developed in `CaRNAval` to extract RINs. This algorithm is also graph based and relies on a greedy approach: it generates seeds (basically minimal common subgraphs) and tries to extend them step by step. The decision of limiting RINs to exactly two SSEs was legitimate as it is a property of known motifs `CaRNAval` was looking to extract (such as A-minors for instance) but it was also a necessary limitation of the search space given the performances of the greedy algorithm. On the contrary, our method does not need such limitation: we can work with any number of SSEs and are thus able to extract more structures, starting with this *RIN^ab^* class.

While being able to study the *RIN^ab^* class was our initial motivation for developing this new method, it quickly appeared that its performances allow the extraction of even broader classes. Indeed, our new method is able to extract *RIN^ab^* more than 50 times faster than `CaRNAval` was extracting *RIN^abc^* (despite working on a generalization of the initial problem). As a consequence, we were able to relax rule *b* (having 2 or more long-range interactions) which, similarly to rule *c*, was a property of known motifs but still partially motivated by the reduction of the search space it induces, and extended our study to the *RIN^a^* class.

#### 3.2.1 Comparison of the *RIN^abc^* and *RIN^ab^* classes

The results presented in `CaRNAval` consist in 331 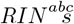 for a total of 6056 occurrences (observation of a RIN in the dataset). From the same dataset, our new method has extracted 557 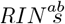 for a total of 7709 occurrences. Amongst the 337 *RIN^abc^ s*, 243 are isomorphic to a *RIN^ab^*. Of the remaining 94 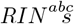, 88 are found inside larger 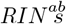, i.e. the canonical graph of the *RIN^abc^* is a subgraph of the canonical graph of at least one *RIN^ab^*. To put it differently, those 88 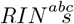 are still captured but are always found inside “larger contexts” that could not be perceived before because of the limitation on the number of SSEs. Now that we relaxed rule *c*, the “larger contexts” are now captured inside 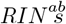 that “assimilated” those 88 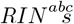. We elaborate further on the question of the SSEs in subsection 3.2.3. For the same reason, the numbers of observations of the 243 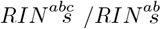 common to both versions have changed for 81 of them (+4 observations on average). All the signal captured by the original version of CaRNAval is present in the new results: all observations of any of those 331 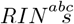 are covered by at least one observation of a *RIN^ab^*.

Please note that `CaRNAval` actually presents 337 structures. However, 4 of them are actually invalid and should not have passed the filters, their absence in our results actually validates our method. The 2 last missing structures are linked to a change in our policies: both have only 2 observations with both observations inside a single RNA chain. We now consider those structures as not recurrent, thus it is normal for our method not to find them. Please also note that we are able to test a graph against itself but choose not to do so.

#### 3.2.2 Network of *RIN^ab^*

Let us now compare the *RIN^abc^* network with the 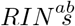 (cf. subsection 2.5.3). The network formed by the 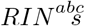 consists in 3 main connected components and named after a characteristic motif they contain. They are the Pseudoknot mesh, the A-minor mesh and the Trans W-C/H mesh, respectively containing 59, 196 and 22 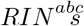. The remaining 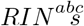 are shared between 25 other components, of size going from 1 to 4.

In contrast, the network of *RIN^ab^* only has 16 components compared to the 28 of the *RIN^abc^* network. It suggests that the newly found 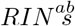 connect components of the *RIN^abc^* network together. This claim is supported by the fact that, in the network of *RIN^ab^*, the Pseudoknot and A-minor meshes have merged into a single one containing 482 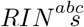. This new giant mesh contains all the elements in the two main meshes presented in `CaRNAval` plus 230 extra 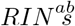. The Trans W-C/H mesh remains disconnected and gains 16 elements for a total of 38 *RIN^ab^*.

#### 3.2.3 *RIN^abc^* and SSEs

The main artificial constraint on 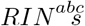 was their restriction to exactly two SSEs. While biologically justifiable, it allowed to strongly constrain the problem making the previous method computable on a large server. By removing this constraint, we observe 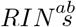 containing varied numbers of SSEs. We show in Fig 7 the distribution of SSEs in the 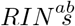 and of their occurrences.

**Figure 7:**
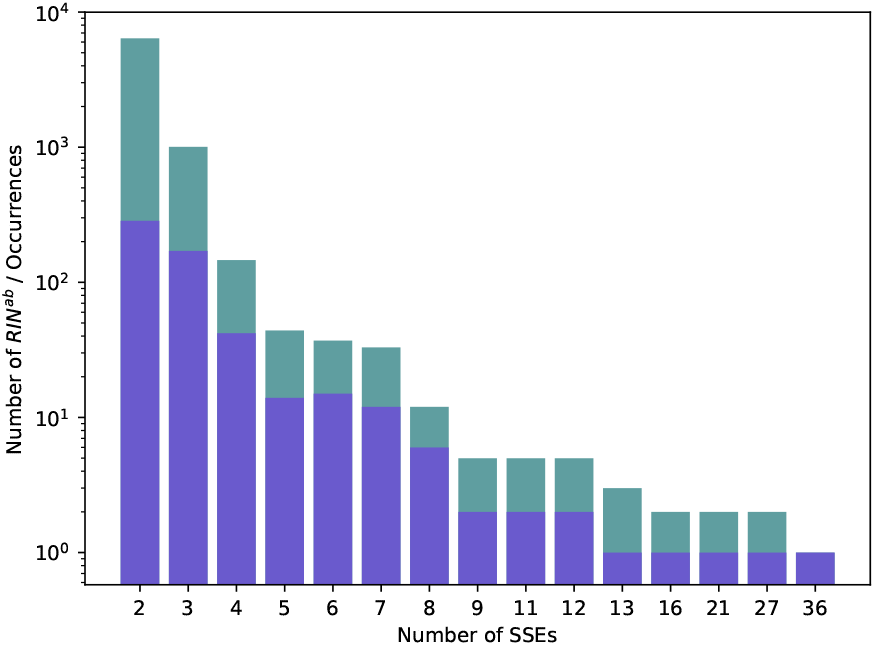
Distribution of 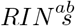 (in blue) and all their occurrences (in green) over the different numbers of SSEs.

Moreover, the different occurrences of the same *RIN^ab^* may contain different numbers of SSEs. We show in Table. 3 that it is globally not the case. Out of the 557 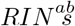, 435 had all of their occurrences span the same number of SSEs. There are 116 that can be over two different number of SSEs, and only 6 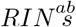 have their occurrences cover three different number of SSEs.

**Table 3:**
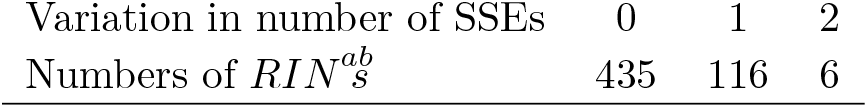
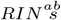 and variation on SSEs span. For each *RIN^ab^* we compute how the number of SSEs covered varies between the occurrences. A value of 0 means that all occurrences are over the same number of SSEs while ±1 (resp. ±2) means that the *RIN^ab^* can span two different number of SSEs (resp. three).

#### 3.2.4 Large 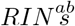

While the largest *RIN^abc^* has 26 nodes, a *RIN^ab^* can potentially encompass an entire molecule. There are 64 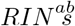 with more than 26 nodes, amongst them 4 have above 100 nodes, the largest *RIN^ab^* containing 293 nucleotides. Those new giants are found in structures of ribosomal subunits. The existence of those 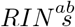 shows that the dataset we are using contains extremely similar structures. The RNA3DHub non-redundant RNAs can still share a considerable portions of their geometry, on up to 293 connexe nucleotides. As a consequence, we might have to update our method, either by modifying our definition of *RIN^ab^* to limit their size or by adding an additional screening to the dataset.

### 3.3 RIN^a^

In the previous section we created the *RIN^ab^* class as a generalization of the *RIN^abc^* class. A natural way to relax even further the problem is to remove the constraint of having 2 or more long range interactions. We call *RIN^a^* the class obtained from *RIN^ab^* by removing rule *b* (cf. definition of the classes in 3.2). While this modification is trivial to implement, the search space increases drastically.

#### 3.3.1 Collection of *RIN^a^*

Our method finds 920 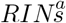 for a total of 12 239 occurrences. All 557 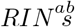 have their canonical graph isomorphic to the canonical graph of a *RIN^a^*. The *RIN^abc^* to *RIN^ab^* transition was done by allowing more than 2 SSEs, which opened the possibility of finding new larger “including” structures. In contrast removing the constraint on the number of long range interactions does not.

We show in Fig. 8 the distribution of the 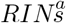 and of their occurrences depending on the number of long range interactions they have. Amongst the remaining 363 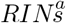, 222 contain no long range interaction and 141 have exactly 1. Those represent 39% of the 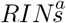 and 37% of the occurrences.

**Figure 8:**
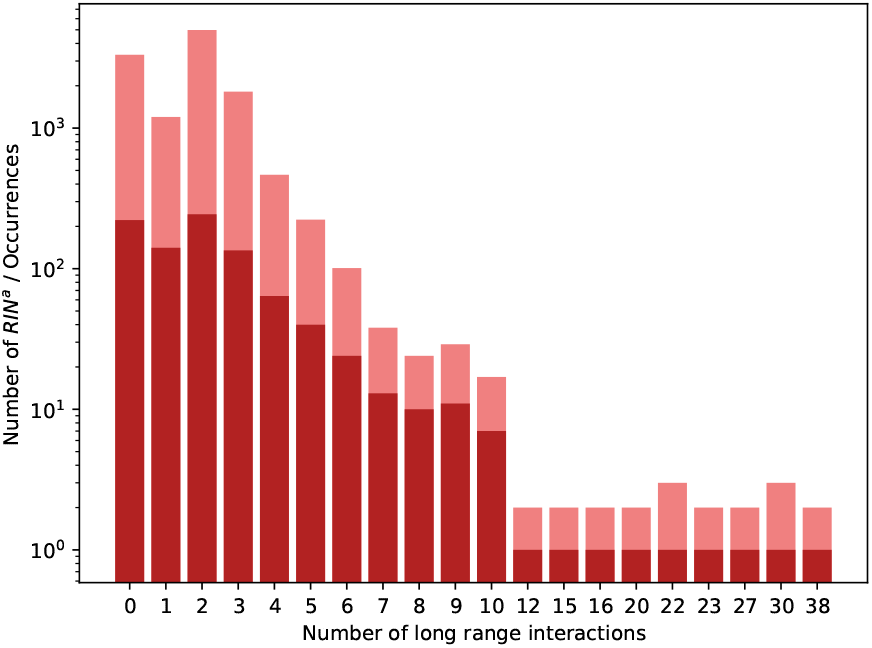
Distribution of 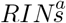 (in red) and all their occurrences (in rose) over the different numbers of long range interactions they contain.

In Fig. 9 we show the distribution of the number of SSEs that are covered by the 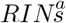. Compared to previously, most 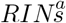 span two SSEs. This shift from the previous, more constrained, results is due to the 222 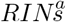 with no long range interactions.

**Figure 9:**
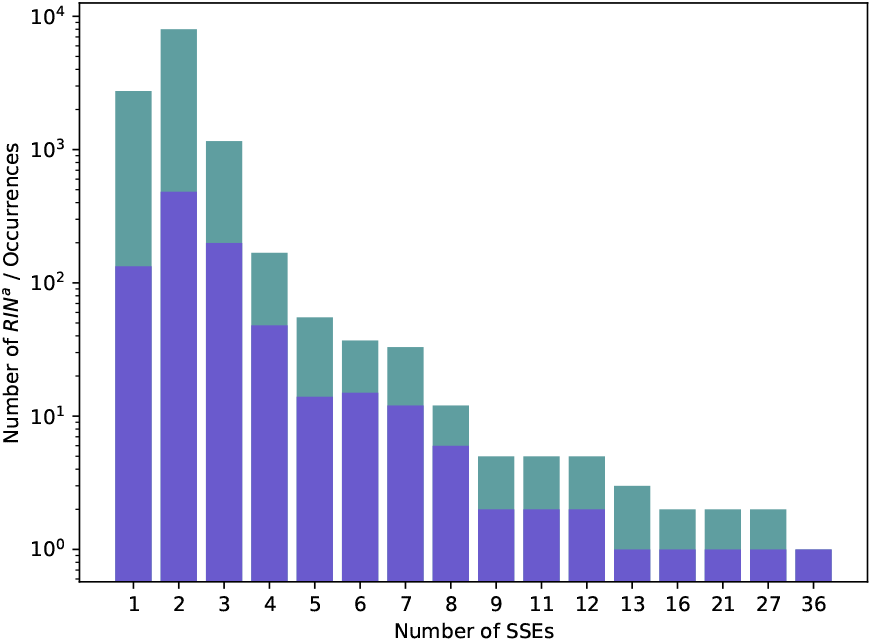
Distribution of 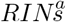 (in blue) and all their occurrences (in green) over the different numbers of SSEs.

As previously, for any given *RIN^a^* the occurrences span a consistent number of SSEs. As we show in Table 4, the same trend as for *RIN^ab^* is followed.

**Table 4:** Variation in the number of SSEs over the occurrences of the same *RIN^a^* (Cf. Table 3). Those numbers show that the variation in the number of SSEs amongst the occurrences of a given *RIN^a^* is both uncommon and limited, even more than with *RIN^ab^*, albeit slightly (82% of 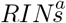 with no variation vs 78% of 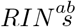).

| Variation in number of SSEs | 0 | $\pm 1$ | $\pm 2$ |
| --- | --- | --- | --- |
| Numbers of $RIN^a$ | 754 | 159 | 7 |

#### 3.3.2 Network of *RIN^a^*

The addition of the new structures to the *RIN^ab^* network connects almost all the nodesof the network. Indeed 888 of the 920 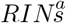 are inside a single giant component. This component gathers not only the Pseudoknot and the A-minor meshes of the *RIN^abc^* network (like the main component of the *RIN^ab^* network did), but also the Trans W-C/H mesh. Of the remaining 32 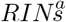 that are not in this component, 23 are singletons, and it remains 6 mini components. In summary, the *RIN^a^* network shows that the *RIN^a^* class forms a unified and nearly totally connected landscape of structures.

#### 3.3.3 Performances

Reproducing the `CaRNAval` dataset we tested the validity of our method and its performances. As all the RINs found and all their occurrences were present in the collection of 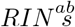, it shows that our method captured strictly more signal than the previous one. In term of performances, the runtime dropped from around _~_330 hours to _~_200 minutes (both are total runtime over the same 20 cores, for the same dataset), despite solving a more general problem. Even relaxing the problem to a maximum by computing the *RIN^a^* class still only took 19 hours in total.

### 3.4 Applications to RNA 3D module-based RNA structure prediction

As described earlier, the rule system of our method allows for a wide range of targets for extraction. We illustrated this with a set of related structure classes (*RIN^abc^,RIN^ab^* and *RIN^a^*. In addition to those classes, we worked on another one linked to the RNA 3D structure prediction problem and *RNA 3D modules. RNA 3D modules* are small RNA substructures involved in structural organization and ligand binding processes. They can be defined with rules similar to the ones describing RINs, with two major differences. First, RNA modules do not need to include long range interactions, and many of the well characterized modules are entirely local, namely the kink-turn and g-bulged modules. Second, unlike RINs, RNA modules are defined by both their structure and sequence profile rather than exclusively the former.

RNA 3D modules can be leveraged in the prediction of a full 3D structure. The fragment-based method implemented by Parisien and Major in MC-Sym[17] constructs a full 3D structure from an augmented secondary structure by mapping the components of this secondary structure to a database of 3D structure fragments. The prediction of 3D modules has been shown to improve this class of methods by providing more informative fragments, namely in RNA-MoIP[21]. Further progress has since been made in this direction with recent improvements in RNA 3D modules identification in sequences[31][25].

The main limitation of this type of method remains the difficulty of assembling a strong dataset of modules. RNA modules are typically identified by searching RNA 3D structures for recurrent subgraphs, a task to which `CaRNAval` should be able to contribute. Unfortunately, as of now, no fragment-based method has been able to integrate long-range modules into a 3D structure prediction pipeline, and the published version of caRNAval cannot be applied to the discovery of common subgraphs without long range interactions as its execution time would explode.

However, the modularity of the methods previously presented (RNA 3D modules basically form the 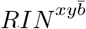, cf. section 3.2, with 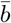 being the constraint of having no long-range interactions), as well as the improved complexity allow for the tackling of this problem. The implementation of those methods constitutes the first software able to discover both long-range and local RNA modules and as such, a significant step towards more accurate fragment-based prediction of 3D structure from sequence.

## 4 Conclusion

In this paper we present a novel method that can find arbitrarily large recurrent interaction networks (RINs) between two RNA structures, represented as graphs. Our method is based on three novel graph matching algorithms (isomorphism, subgraph and maximal common subgraph algorithms) that leverage the proper edge coloring property we exhibited in RNA structures represented as graphs. Those novel algorithms improve drastically on previous methods, (notably being a hundred time faster than the most comparable other method: `CaRNAval`), and allow for the first time to identify modules arbitrarily large. Moreover our method distinguishes between the rule system used to define the structures to extract and the graph matching algorithms used to extract them. As a consequence it is able to extract a broad range of structures and to easily switch between targets to extract. The gain in efficiency allows to relax the constraints and search for broad classes of RINs like *RIN^a^*, which can span any number of SSEs, and have any number of long range interaction, even none.

In `CaRNAval` the network of found modules had three clear main components. We show that the network of found 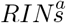 is a massive components linking together more that 95% of the recurrent structures together. This can be key to understand how those structural features emerged and were propagated, and to improve design of artificial RNAs.

## Footnotes

1 We will be using *class* to refer to subsets of structural elements from now on as the relations between subsets are similar to the ones between the classes of a class-oriented langage.

