## Supplementary Material for "Finding recurrent RNA structural networks with fast maximal common subgraphs of edge-colored graphs"

**Antoine Soulé [1,2], Vladimir Reinharz [3], Roman Sarrazin-Gendron [1], Alain Denise [4,5,*] and Jérôme Waldispühl [1,*]**

[1]School of Computer Science, McGill University, Montreal, Canada

[2]LiX, École Polytechnique, Paris, France

[3]Department of Computer Science, Université du Québec à Montréal, Montreal, Canada

[4]Université Paris-Saclay, CNRS, Laboratoire de recherche en informatique, Palaiseau, France

[5] Université Paris-Saclay, CEA, CNRS, Institute for Integrative Biology of the Cell (I2BC), Gif-sur-Yvette, France

$*$ Corresponding Authors

April 2020

### Chapter 1

### Algorithms for efficient graph matching of edge-colored graphs

#### 1.1 General definitions

The algorithms presented here take two connected and undirected graphs $G = \{V_G, E_G\}, H = \{V_H, E_H\}$ as input, where $V_G$ and $V_H$ represent the sets of vertices/nodes and $E_G$ and $E_H$ the sets of edges.

The edges in $E$ (whether $E_G$ or $E_H$) are colored and the set of all colors $L$ has a finite size. $E_l$ is defined such as $\forall e \in E_l$ the color of $e$ is $l$. As a consequence $\forall l, E_l \subset E$ and $E = (\bigcup_{l \in L} E_l)$.

A node $v'$ is in the *neighbourhood* of a node $v$ *iff* $\exists (v, v') \in E$. The neighbourhood of a node $v$ is denoted $N(v)$. We extend this notion to every color $l \in L : v' \in N_l(v) \iff \exists (v, v') \in E_l$.

$G$ and $H$ are equipped with a *proper edge coloring* : the colors of edges incident to the same vertex are different. To put it differently : $\forall v \in V, \forall l \in L, |N_l(v)| \leq 1$, each node is involved in at most one edge of a given color.

#### 1.2 Isomorphism

##### 1.2.1 Definition of the problem

$G = \{V_G, E_G\}$ and $H = \{V_H, E_H\}$ are isomorphic *iff* $\exists$ a bijection $b$ from $V_G$ to $V_H$ that respects the edges and their colors, namely $\forall g_1, g_2 \in V_G, \forall h_1, h_2 \in V_H$ if $b(g_1) = h_1 \land b(g_2) = h_2$, then $\{g_1, g_2\} \in E_{l,G} \iff \{h_1, h_2\} \in E_{l,H}$.

The problem is to determine if two given connected and undirected graphs $G$ and $H$ are isomorphic.

##### 1.2.2 Algorithm

*Note 1 :* We assume that the graphs $G$ and $H$ are *potentially isomorphic i.e.* :
1. $|V_G| = |V_H|$
2. $\forall l \in L, |E_{l,G}| = |E_{l,H}|$

Those properties are necessary for those graphs to be isomorphic, can be checked in linear time and assuming them simplifies the description of the algorithm.

*Note 2 :* We use *mapping* objects to represent the set of matched nodes. A *mapping* can be seen here as a simple collection of pairs of nodes $(g, h)$ with $g \in V_G$ and $h \in V_H$. Given such a *mapping* $m$, we will also denote $g \in m$ (resp $h \in m$) if there exists a pair $(g, h) \in m$.

*Intuition of the strategy :* The algorithm aims at extending partial matchings of $G$ and $H$ until they are complete. For two nodes $g$ and $h$ matched, the *proper edge coloring* guarantees that there will be only one way of matching their respective neighbourhoods thus making the extension of a matching linear in the size of the graphs.
If the neighbourhoods cannot be matched at any point of the extension, the current matching cannot be completed and thus is discarded. If no matching can be extended until completion (i.e. until the mapping under construction reach the size of $G$ and $H$ thus making it a bijection), the two graphs are not isomorphic.

Our algorithm solving the *graph isomorphism* problem comes in three parts.

- `Launcher`$(G, H)$ which will consider all eligible starting positions from graphs $G$ and $H$, create a mapping $m$ from it and call `Extender`$(m)$.
- `Extender` which will try to extend the matching $m$ as much as possible.
- `EligibleNeighbours`$(m, g, h)$ is used by `Extender` to obtain the pairs of nodes from $G$ and $H$ which are in the neighbourhood of the pair `Extender` is currently processing and that are eligible to be added to the mapping.

---

**Algorithm 231:** `Launcher`$(G, H)$

---

**Data:** Two connected and undirected graphs $G$ and $H$
**Result:** *True* if $G$ and $H$ are isomorphic, *False* otherwise.

$l \leftarrow argmin_{\ l' \in L}\{|E_{l',G}|\}$
**for** $e_G \in E_{l,G}, e_H \in E_{l,H}$ **do**
  $\{g_1, g_2\} \leftarrow e_G$
  $\{h_1, h_2\} \leftarrow e_H$
  $m \leftarrow \{(g_1, h_1)\} \cup \{(g_2, h_2)\}$
  **if** $Extender(m)$ **then**
    **return** $True$
  $m \leftarrow \{(g_1, h_2)\} \cup \{(g_2, h_1)\}$
  **if** $Extender(m)$ **then**
    **return** $True$
**end**
**return** $False$

---

**Algorithm 232:** `Extender`($m$)

**Data:** A Mapping $m$
**Result:** $True$ if $m$ has been extended to completion, $False$ otherwise

```
queue f ← ∅
for (g, h) ∈ m do
    f.add((g, h))
end
while f ≠ ∅ do
    (g, h) ← f.pop()
    if EligibleNeighbours(m, g, h) == ⊥ then
        return False
    for (g', h') ∈ EligibleNeighbours(m, g, h) do
        f.add((g', h'))
        m ← m ∪ {(g', h')}
    end
end
return True
```

**Algorithm 233:** `EligibleNeighbours`($m, g, h$)

**Data:** A mapping $m$, mapped nodes $g$ and $h$
**Result:** The set $s$ of pairs of mappable neighbours of $g$ and $h$

```
c ← ∅
for l ∈ L do
    if |N_l(g)| ≠ |N_l(h)| then
        return ⊥
    if |N_l(g)| > 0 then
        g' ← N_l(g)
        h' ← N_l(h)
        if (g', h') ∉ m then
            if g' ∈ m ∨ h' ∈ m then
                return ⊥
            c ← c ∪ {(g', h')}
end
return c
```

##### 1.2.3 Complexity

The proper edge coloring trivially implies that the maximal degree of $G$ and $H$ is smaller than $|L|$ thus their sizes are linear in their order. It also implies that $max_{l \in L}(|E_l|) \leq |V|/2$.

The extension is linear in the size of the graphs and so is linear in their order (as the degree of the vertices is bounded by the number of colors which is finite). The number of starting points (cf. Alg. 231 l.2) is bounded by $2 \times (|V|/2)^2$. As a consequence, the complexity of this algorithm is $\mathcal{O}(n^3)$ with $n = |V|$ (since we assumed the two graphs have the same order).

*Comparison with existing algorithms* Even if it has not been proven NP-hard, the isomorphism problem is NP in the general case [1, 2, 3]. However, polynomial algorithms have been found for several classes of graphs [1, 3]. None of those classes strictly match the graphs we are working with but the degree of those graph is bounded by the number of different colors and thus those graphs are of bounded degree. An upper bound for the complexity of the isomorphism problem for graphs of bounded degree $k$ has been proposed by Luks [4] : $\mathcal{O}(n^{c\ k\ log(k)})$ with integer $c > 1$. Our algorithm performs better for all $k > 4$, even under the optimistic assumption that $c = 2$. As $k = |L|$, it thus gives our algorithm the edge when there are at least 4 different colors and this advantage increases greatly with the number of colors as the complexity of our algorithm is not impacted by it.

#### 1.3 Subgraph isomorphism

##### 1.3.1 Definition of the problem

A graph $G = \{V_G, E_G\}$ is a *subgraph* of graph $H = \{V_H, E_H\}$ *iff* there exists at least one injection $i$ from $V_G$ to $V_H$ such that: $\forall g, g' \in V_G$, $\{g, g'\} \in E_{l,G} \implies \{i(g), i(g')\} \in E_{l,H}$

The problem is, given two connected and undirected graphs $G$ and $H$, to determine whether $G$ is a subgraph of $H$.

##### 1.3.2 Algorithm

*Note 1 :* We assume $G$ is a *potential subgraph* of $H$ *i.e.*
1. $|V_G| \leq |V_H|$
2. $\forall l \in L, |E_{l,G}| \leq |E_{l,H}|$

Those properties are necessary for $G$ to be a *subgraph* of $H$, can be checked in linear time and assuming them simplifies the description of the algorithm.

*Intuition of the strategy :* This algorithm is an evolution of the isomorphism test, the only difference being the stop condition. Indeed, it only requires the neighbourhood of $g$ to be matched to the neighbourhood of $h$ and not the converse.
Our algorithm solving the *subgraph isomorphism* problem follows the same architecture as the one solving *graph isomorphism* problem. As a consequence, it uses the same methods. The difference with the previous algorithm lays in the conditions inside the methods.

**Algorithm 241:** `Launcher`$(G, H)$

**Data:** Two connected and undirected graphs $G$ and $H$
**Result:** $True$ if $G$ is a subgraph of $H$, $False$ otherwise.

$l \leftarrow argmin_{\ l' \in L}\{|E_{l',G}| \times |E_{l',H}| \text{ s.t. } |E_{l',G}| > 0\}$
**for** $e_G \in E_{l,G}, e_H \in E_{l,H}$ **do**
  $\{g_1, g_2\} \leftarrow e_G$
  $\{h_1, h_2\} \leftarrow e_H$
  $m \leftarrow \{(g_1, h_1)\} \cup \{(g_2, h_2)\}$
  **if** $Extender(m)$ **then**
    **return** $True$
  $m \leftarrow \{(g_1, h_2)\} \cup \{(g_2, h_1)\}$
  **if** $Extender(m)$ **then**
    **return** $True$
**end**
**return** $False$

**Algorithm 242:** `Extender`$(m)$

**Data:** A mapping $m$
**Result:** $True$ if $m$ has been extended to completion, $False$ otherwise

queue $f \leftarrow \emptyset$
**for** $(g, h) \in m$ **do**
  $f.add((g, h))$
**end**
**while** $f \neq \emptyset$ **do**
  $(g, h) = f.pop()$
  **if** $EligibleNeighbours(m, g, h) == \perp$ **then**
    **return** $False$
  **for** $(g', h') \in EligibleNeighbours(m, g, h)$ **do**
    $f.add((g', h'))$
    $m \leftarrow m \cup \{(g', h')\}$
  **end**
**end**
**return** $True$

**Algorithm 243:** `EligibleNeighbours`$(m, g, h)$

**Data:** A mapping $m$, mapped nodes $g$ and $h$

**Result:** The set $s$ of pairs of mappable neighbours of $g$ and $h$

```
c ← ∅
for l ∈ L do
    if |N_l(g)| > |N_l(h)| then
        return ⊥
    if |N_l(g)| > 0 then
        g' ← N_l(g)
        h' ← N_l(h)
        if (g', h') ∉ m then
            if g' ∈ m ∨ h' ∈ m then
                return ⊥
            c ← c ∪ {(g', h')}
end
return c
```

##### 1.3.3 Complexity

The only difference with the previous algorithm is that $|V_G| \leq |V_H|$, thus the complexity of this algorithm is $\mathcal{O}(n^3)$ with $n = |V_H|$.

*Comparison with existing algorithms* The subgraph problem is NP-hard in the general case. The closest graph class covering our graphs are the bounded degree graphs. The complexity of the best algorithm solving the isomorphism problem for graphs of bounded degree $k$ is in $\mathcal{O}(n^{k*log(k)})$.

#### 1.4 Maximal Common Subgraph

##### 1.4.1 Maximal Common Subgraph Definitions

**Definition 1. Common Subgraph (csg) :** Given two graphs $G$,$H$, a graph $S = (V_S, E_S)$ is a *common subgraph* ($csg$) of $G$ and $H$ if it is a subgraph of $G$ and a subgraph of $H$.

**Definition 2. Equality of csg :** Let us consider $S = (V_S, E_S)$ and $S' = (V_{S'}, E_{S'})$ two $csg$ of $G$ and $H$, $i_G$ (resp. $i_H$,$i'_G$,$i'_H$) the injection from $V_S$ (resp. $V_S$,$V_{S'}$,$V_{S'}$) to $V_G$ (resp. $V_H$,$V_G$,$V_H$). $S = S'$ iff $S$ and $S'$ are isomorphic and the corresponding bijection $b : V_S \rightarrow V_{S'}$ satisfies the additional conditions: $i_G = i'_G \circ b$ and $i_H = i'_H \circ b$.

**Definition 3. Maximal Common Subgraph (mcsg) :** A $csg$ $S$ of $G$ and $H$ is *maximal* ($mcsg$) iff for all $S'$ subgraph of $G$ and $H$, $S \subset S' \implies S = S'$ (in the sense of definition 2).

**Maximal Common Subgraph Problem :** The *Maximal Common Subgraph problem* usually refers to finding a csg $S = (V_S, E_S)$ of graphs $G$,$H$ such that for all csg $S' = (V_{S'}, E_{S'})$ of graphs $G$,$H$, $|V_S| \geq |V_{S'}|$.
However the algorithm we propose produces instead the set $MCSG_{G,H}$ of all mcsg of graphs $G$ and $H$. The usual problem of finding the biggest csg of graphs $G$,$H$ is obviously reducible to the one we are solving here as the biggest element of $MCSG_{G,H}$ is the biggest csg of graphs $G$,$H$.

##### 1.4.2 Algorithm & Associated Definitions

This algorithm, while being related to the two previous ones, introduces major evolutions in order to manage a crucial difference with the two previous problems: a discrepancy no longer causes a termination but instead open another path to explore. We provide here the key elements needed to understand the algorithms. A more complete and formal description is provided in section 1.4.3.

**Starting Point :** A *starting point* $\alpha$ is defined for two graphs $G = \{V_G, E_G\}$ and $H\{V_H, E_H\}$ as $\alpha = \big((g_1, h_1), (g_2, h_2)\big)$ with $g_1, g_2 \in V_G, h_1, h_2 \in V_H, \{g_1, g_2\} \in V_{l,G}, \{h_1, h_2\} \in V_{l,H}$. In other words, the set of starting points is exactly the set of common subgraphs of size two.

**Division of the search space** The set of maximal common subgraphs $MCSG_{G,H}$ between two graphs $G$ and $H$ is produced by the algorithm as $\bigcup_\alpha mcsg_{G,H,\alpha} = MCSG_{G,H}$ where $mcsg_{G,H,\alpha} \subset MCSG_{G,H}$ corresponds to the sets of maximal common subgraphs produced from the starting point $\alpha$. $mcsg_{G,H,\alpha}$ is also the set of all maximal common subgraphs containing $\alpha$. As a consequence, we will often work with a specific starting point $\alpha$ and consider the set $mcsg_{G,H,\alpha}$ rather than the whole $MCSG_{G,H}$ in section 1.4.3.

**Conflicts** We have to introduce the notion of conflict that was implicite in both the isomorphism problem and the subgraph problem. Indeed, any discrepancy in those two problems implies that the extension cannot lead to a solution and thus provoked termination ($\perp$). However, a discrepancy now suggests the existence of another way to map $G$ and $H$ and so need to be considered. As a consequence we shall introduce the notions of *conflict* and *branch.*

We call *conflict* the event in which a pair $(g, h)$ considered for addition to the mapping cannot be added because the mapping already contains either a pair $(g, h')$ or a pair $(g', h)$ or both (with $g \neq g'$ and $h \neq h'$ obviously). As a consequence, we describe a conflict as a tuple $((g, h), P)$ where $(g, h)$ is the pair that cannot be added and $P$ the list of pairs that prevents its addition and already are in the mapping ($P$ contains either one or two pairs depending on the number of pairs $(g, h)$ is in conflict with).

A conflict suggests the existence of a different mcsg than the one currently being produced and thus the need to launch another exploration with additional constraints so it could produce it by prohibiting the addition of the pairs in $P$. We call *branch* an exploration with a given set of constraints.

**Branches and binary words** As the exploration is deterministic and is only constrained by interdictions of forming specifics pairs, the behaviour of any branch $B \in mcsg_{G,H,\alpha}$ can be described with a binary word $\beta$, of which each element/letter $a$ cor-

responds to the interdiction of matching the two nodes $g \in V_G$ and $h \in V_H$ in the pair $a = (g, h)$. Several binary words can describe the same branch but each branch admits a single minimal (lengthwise) binary word containing only the pairs which interdiction actually impacts the extension. We will denote a branch by a tuple $\{\alpha, \beta\}$.
Note that the first branch explored is denoted by the empty word $\epsilon$ as there is no interdiction.

**Extension-Induced Order** As the extension process is deterministic, the production of a mapping $m$ with the starting point $\alpha$ (i.e. $m \in mcsg_{G,H,\alpha}$) in the branch $\beta$ implies an order on all pairs $(g, h)$ considered during the extension, whether it has been accepted and added to the mapping or rejected. We denote $o_{\alpha,\beta}$ the order induced by the exploration of branch $\beta$ from starting point $\alpha$. Each order is partial if we consider $mcsg_{G,H,\alpha}$ but we will only need to compare elements considered during the same exploration and thus covered by the corresponding partial order. We represent $o_{\alpha,\beta}$ as an ordered list of pairs $(g, h) \in \{V_G \times V_H\}$ considered during the exploration of branch $\beta$. In addition to the order, we also preserve for each pair $(g, h)$ the decision the algorithm has made, represented by a character in $\{A|F|C\}$. Indeed, the pair $(g, h)$ could either have been $A$dded to the mapping, rejected because it was in $\beta$ and thus $F$orbidden or rejected because the mapping already contained a pair $(g, h')$ or $(g', h)$ or both, generating a $C$onflict.

We will denote $a <_{o_{\alpha,\beta}} b$ the comparison of the positions of pairs $a$ and $b$ in the order $o_{\alpha,\beta}$.

**Mapping** We use *mapping* objects to represent the set of matched nodes and store several informations. The *mapping*s of this section have the following properties :

- a *mapping* is a collection of pairs of node $(g, h)$ with $g \in V_G$ and $h \in V_H$
- a binary word $\beta$ is associated to the *mapping* $m$ and represents the specificities of the branch the extension of mapping $m$ corresponds to
- the starting point $\alpha = ((g_1, h_1), (g_2, h_2))$ with $g_1, g_2 \in V_G, h_1, h_2 \in V_H, \{g_1, g_2\} \in V_{l,G}, \{h_1, h_2\} \in V_{l,H}$. $\alpha$ is the first addition to $m$ and cannot be challenged

- the order $o_{\alpha,\beta}$ in which the pairs of nodes have been considered during the exploration

For convenience, $m[x]$ denotes the node $x$ is mapped to in the mapping $m$ (i.e. for any pair $(g,h) \in m$ with $g \in G$ and $h \in H$, $m[g] = h$ and $m[h] = g$). Given such a *mapping* $m$, we will also denote $g \in m$ (resp $h \in m$) if there exists a pair $(g,h) \in m$.

**Conflict management** As mentioned before, a conflict suggests the existence of a maximal common subgraph that cannot be produced by the current exploration and thus the need to create a new branch potentially able to produce this new maximal common subgraph.

To describe the process of creating a new branch, let us consider a generic case where an exploration from starting point $\alpha$ associated with binary word $\beta$, that produced a maximal common subgraph, discovered a conflict $((g,h),P)$. The new branch will use the same starting point but a different binary word that we will denote $\beta'$. Recall that those binary words represent pairs of nodes that we are not allowed to form. We know that, for the new branch to be able to form $(g,h)$, the pairs in $P$ need to be in $\beta'$ however we have to determine which part of $\beta$ must be transferred in $\beta'$.

Since the branch $(\alpha,\beta)$ has been explored, we have that the orders $o_{\alpha,\beta}$ compares $(g,h)$, all pairs of $\beta$ and all pairs in $P$. We consider that all pairs considered by the exploration before it reaches $(g,h)$ may be necessary for it to reach $(g,h)$. As a consequence $\beta'$ will contain all pairs $(g',h')$ such that $o_{\alpha,\beta}[(g',h')] < o_{\alpha,\beta}[(g,h)]$ plus the pairs in $P$.

If the exploration of branch $(\alpha,\beta)$ encountered several conflicts, this process is repeated for each of them.

If the exploration of branch $(\alpha,\beta)$ did not produce a maximal common subgraph (we will discuss the reasons for such scenario in section 1.4.4), no new branches are created from branch $(\alpha,\beta)$ (cf. 1.4.2).

As several branches may lead to the creation of the same new branch $(\alpha,\beta')$, we keep a record of which tuples $(\alpha,\beta')$ have been created and explored so we don't explore twice the very same branch as it would be a waste.

For instance and as displayed in fig. 1.1, the exploration of the branch $\{\alpha,\beta_1 = \epsilon\}$ discovers two conflicts implying mapped pairs $a$ and $b$. The algorithm thus creates two new branches $\{\alpha, \beta_2 = a\}$ and $\{\alpha, \beta_3 = b\}$. However, the exploration of branch $\{\alpha, \beta_2 = a\}$ leads to the discovery of conflict $(c, \{c'\})$ with $c > a$ and thus the creation of a branch $\{\alpha, \beta_3 = ac\}$ to be explored. If $c < a$ the branch created would have been $\{\alpha, \beta_3 = c\}$. *etc,...*

**Test of maximality** The algorithm detailed before may output csg that are not maximal as we briefly mentioned before. That is why we add a trimming step afterwards which we will describe after the study of this algorithm in section 1.4.4.

**Edge maximality** By definition, maximal common subgraphs are also maximal in terms of edges (i.e. no edge can be added to a maximal common subgraph) thus $\forall S, S'$ maximal common subgraphs of graphs $G$ and $H$, $V_S = V'_S \iff E_S = E'_S \iff S = S'$.

**Algorithms** Our algorithm solving the *maximal common subgraphs* is based on the same core as the two previous one but comes with an additional parts for a total in four parts.

- `Launcher`$(G, H)$ which will consider all eligible starting points $\alpha$ from graphs $G$ and $H$ and call `Branch_Management`$(\alpha)$.
- `Branch_Management` will create a mapping $m_{\alpha,\beta}$ with $\beta = \epsilon$, call `Extender`$(m_{\alpha,\beta})$ and process its output. Processing the output will either return its results or create new branches and thus new calls to `Extender`, etc,...
- `Extender`$(m_{\alpha,\beta})$ will try to extend the matching $m_{\alpha,\beta}$ as much as possible and record the conflict raised in the process.
- `EligibleNeighbours`$(m, g, h)$ is used by `Extender` to obtain the pairs of nodes from $G$ and $H$ which are in the neighbourhood of the pair `Extender` is currently processing and that are eligible to be added to the mapping.

a. Step 1

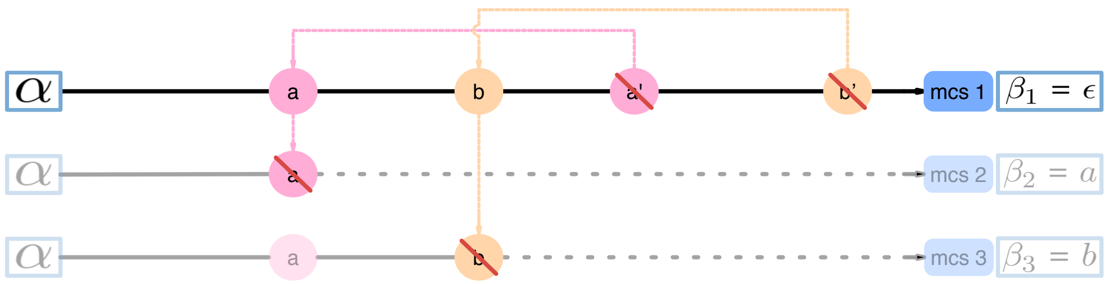

b. Step 2

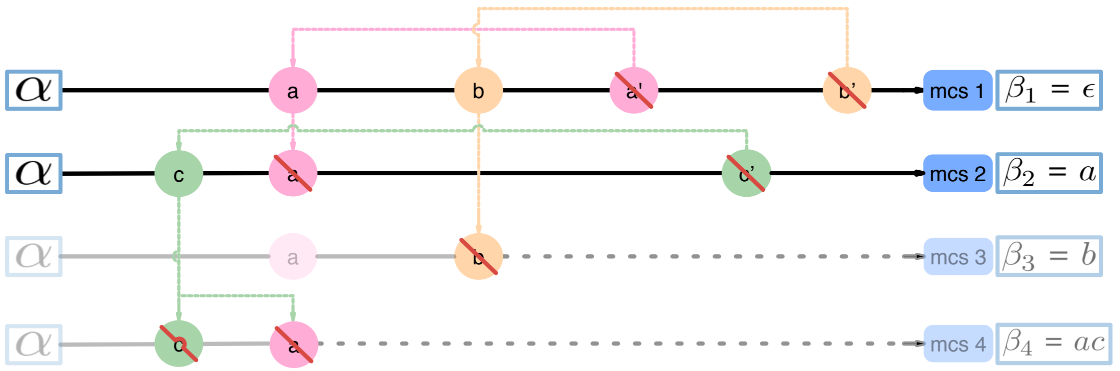

Figure 1.1: **Branching-Backtrack Mechanism:** In this simplified diagram, each row is a linear representation of the non-linear exploration of a branch from the starting point $\alpha$ up to the point where no more pairs are available.

The upper panel shows the result of the exploration of the first branch. As no pairs are forbidden, $\beta_1 = \epsilon$. This exploration encounters two conflicts: $(a, \{a'\})$ in *pink* and $(b, \{b'\})$ in *orange*. Each conflict leads to the creation of a new branch that has yet to be explored. Those branches are represented by the second and third rows respectively. The plain lines represent the part of the exploration that is common with the first branch while the dot lines represent the part that is unknown.

The lower panel panel shows the result of the exploration of the second branch where the formation of pair $a$ is forbidden ($\beta_2 = a$). This exploration encountered a single conflict: $(c, \{c'\})$ in *green* that should lead to the creation of a fourth branch with $\beta_4 = c$. However, since $c'$ has been considered after $a$ was rejected, we assume that forbidding $a$ is also required to form $c'$ thus $\beta_4 = ac$.

**Algorithm 251:** `Launcher`$(G, H)$

**Data:** Two connected and undirected graphs $G$ and $H$

**Result:** The collection of all maximal mappings

```
MCSG_(G,H) ← ∅
for i | 0 ≤ i ≤ λ − 1 do
    for e_G, e_H | e_G ∈ E_{G,i} ∧ e_H ∈ E_{H,i} do
        (g_1, g_2) ← e_G
        (h_1, h_2) ← e_H
        α ← {(g_1, h_1)} ∪ {(g_2, h_2)}
        MCSG_(G,H) ← MCSG_(G,H) ∪ Branch_Management(α)
        α ← {(g_1, h_2)} ∪ {(g_2, h_1)}
        MCSG_(G,H) ← MCSG_(G,H) ∪ Branch_Management(α)
    end
end
return MCSG_(G,H)
```

**Algorithm 252:** `Branch_Management`($\alpha$)

**Data:** A starting point $\alpha$

**Result:** The set of maximal mappings obtained from the given starting point

$mcsg_{(G,H,\alpha)} \leftarrow \varnothing$
queue $branches\_to\_explore \leftarrow \{\epsilon\}$
$created\_branches \leftarrow \{\epsilon\}$
**while** $branches\_to\_explore \neq \varnothing$ **do**
    $\beta \leftarrow branches\_to\_explore.pop()$
    mapping $m \leftarrow$mapping($\alpha$)
    $m.\alpha \leftarrow \alpha$
    $m.\beta \leftarrow \beta$
    $m, C, O \leftarrow Extender(m)$
    `/* The following double loop that creates the new binary words is not optimal */`
    `/* However, the single loop version is more obscure */`
    **if** $m$ *is maximal* **then**
        $mcsg_{(G,H,\alpha)} \leftarrow mcsg_{(G,H,\alpha)} \cup m$
        **for** $\big((g,h),P\big) \in C$ **do**
            $\beta' \leftarrow \epsilon$
            **for** $(g',h') \in \beta$ **do**
                **if** $(g',h') <_O (g,h)$ **then**
                    $\beta' \leftarrow \beta'.(g',h')$
            **end**
            **for** $(g'',h'') \in P$ **do**
                $\beta' \leftarrow \beta'.(g'',h'')$
            **end**
            **if** $\beta' \notin created\_branches$ **then**
                $branches\_to\_explore.put(\beta')$
                $created\_branches \leftarrow created\_branches \cup \beta'$
        **end**
**end**
**return** $mcsg_{(G,H,\alpha)}$

**Algorithm 253:** `Extender`$(m)$

**Data:** A mapping $m$

**Result:** The collection of maximal mappings extended from $m$ and the set $C$ of conflict generated during the extension

$C \leftarrow \emptyset$
$O \leftarrow \emptyset$
queue $f \leftarrow \emptyset$
**for** $(g, h) \in sorted(m.\alpha)$ **do**
    $f.put((g, h))$
**end**
**while** $f \neq \emptyset$ **do**
    $(g, h) \leftarrow f.pop()$
    /* $C$ and $O$ are passed by reference i.e. modifications to them remain */
    **for** $(g', h') \in EligibleNeighbours(m, g, h, C, O)$ **do**
        $f.add((g', h'))$
        $m \leftarrow m \cup \{(g', h')\}$
    **end**
**end**
**return** $m, C, O$

**Algorithm 254:** `EligibleNeighbours`$(m, g, h, C, P)$

**Data:** A mapping $m$, mapped nodes $g$ and $h$, collection of conflicts $C$ and ordered collection $O$

**Result:** The set $s$ of pairs of mappable neighbours of $g$ and $h$

$s \leftarrow \emptyset$
**for** $l \in sort(L)$ **do**
  **if** $|N_l(g)| > 0 \wedge |N_l(h)| > 0$ **then**
    $g' \leftarrow N_l(g)$
    $h' \leftarrow N_l(h)$
    **if** $\overline{(g', h')} \notin m.\beta$ **then**
      **if** $g' \notin m \wedge h' \notin m$ **then**
        $s \leftarrow s \cup \{(g', h')\}$
        $O.add\big(\ ((g', h'), A)\ \big)$
      **else**
        **if** $g' \notin m$ **then**
          $P \leftarrow \big((g', m[h'])\big)$
        **else if** $h' \notin m$ **then**
          $P \leftarrow \big((m[g'], h')\big)$
        **else**
          $P \leftarrow \big((g', m[h']), (m[g'], h')\big)$
        **if** $\forall p \in P,\ p \notin m.\alpha$ **then**
          $C \leftarrow C \biguplus \big((g', h'), P\big)$
        $O.add\big(\ ((g', h'), C)\ \big)$
    **else**
      $O.add\big(\ ((g', h'), F)\ \big)$
**end**
**return** $s$

##### 1.4.3 Proofs & associated definitions

**Exploration binary tree** Even if the algorithm does not use it, the set of all branches in $mcsg_{G,H,\alpha}$ forms an *exploration binary tree* (EBT) that can be produced *a posteriori*. The root of the EBT is the starting point $\alpha$ and each leaf corresponds to the common subgraph produced by a branch. Each path from the root to a leaf describes exactly the exploration of the branch the leaf corresponds to. Each non-leaf node of the EBT corresponds to a pair $(g, h)$. Going right from a non-leaf node representing the pair $(g, h)$ corresponds to the addition of said pair to the current mapping and going left corresponds to its rejection because the pair was in the binary word $\beta$ of the branch currently being explorated. The algorithm to build this tree is described below (cf. algo. 255). As the pair $(g, h)$ may be encountered in several explorations, several non-leaf nodes of the EBT may represent the same pair $(g, h)$. In order to distinguish between those, the non-leaf nodes of the EBT are denoted $\rho(g, h)$ with $(g, h)$ the pair they represent and $\rho \in \{right|left\}^+$ the sequence of decisions made from the root to $\rho(g, h)$.
Even if the EBT is never used by the algorithme and can only be produced *a posteriori* we will be relying on it heavily in this section as it is a convenient way to describe the intrication of the exploration. For this same reason, we will also use expressions such as "when the node $\rho(g, h)$ was created". Since the EBT is created at once, such expressions are not to be taken literally. They actually refer to the first time the algorithm considered the pair $(g, h)$ after the sequence $\rho$ of decisions which is the moment we become sure that the EBT will contain a node $\rho(g, h)$.

**Definition 4.** Consider two csg $S$ and $S'$ of $G$ and $H$, with their corresponding inclusions $i_G : V_S \hookrightarrow V_G$ and $i_H : V_S \hookrightarrow V_H$, $i'_G : V_{S'} \hookrightarrow V_G$ and $i'_H : V_{S'} \hookrightarrow V_H$. The csg $S$ and $S'$ *share a vertex* if there exists $s \in V_S$ and $s' \in V_{S'}$ such that $i_G(s) = i'_G(s')$ and $i_H(s) = i'_H(s')$.

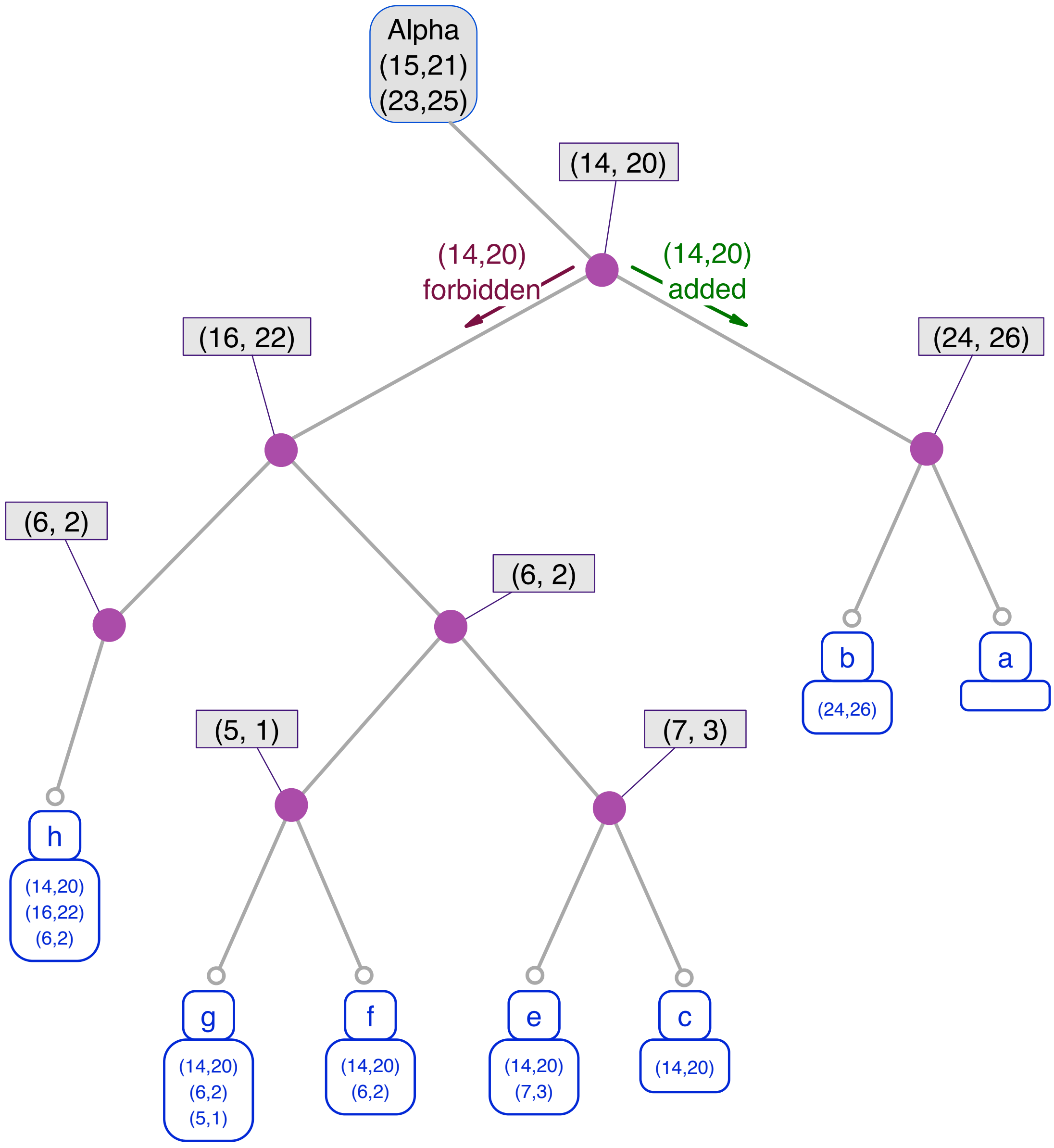

Figure 1.2: **Exploration Binary Tree:** This figure displays the EBT of an actual set of branches generated by the algorithm 255. Each leaf corresponds to a branch denoted from $a$ to $h$ according to the order in which they have been created. The associated binary words $\beta$ are displayed below each branch. For the sake of readability, we collapse all non-leaf nodes without a left son at the exception of the root (i.e. $\alpha$). As a consequence, all the remaining non-leaf nodes correspond to pairs effectively forbidden in at least one binary word $\beta$. Since the default behaviour during the extension of the csg is to accept the pair of nodes the algorithm is currently considering (which corresponds to going right in the EBT), this transformation does not lead to any loss of information as it preserves all meaningful events. For each remaining non-leaf node is displayed the corresponding pair $(g, h)$ (ex: $(14, 20)$). Please note that the tree contains two nodes corresponding to pair $(6, 2)$ which illustrates the need to join the sequence of decisions $\rho$ to designate a node in the EBT.

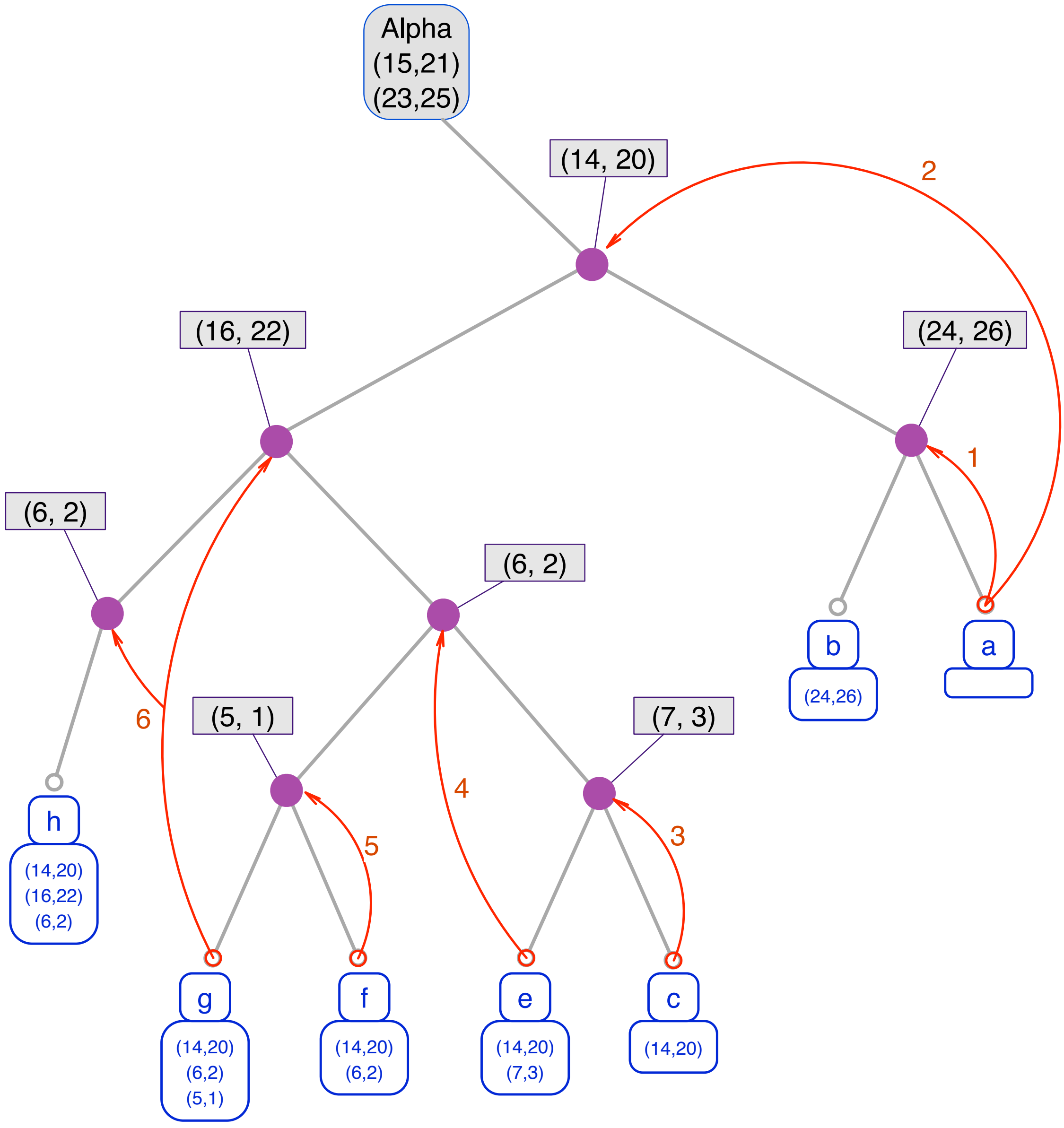

Figure 1.3: **Conflicts and new branches:** This figure complements figure 1.2 with a red overlay showing the relations between conflicts and creations of new branches. For instance, branch $a$ encountered two conflicts. Conflict 1 revealed the necessity of forbidding the pair $(24, 26)$ thus creating branch $b$ and Conflict 2 revealed the necessity of forbidding the pair $(14, 20)$ thus creating branch $c$. Branch $b$ did not encounter any conflict but branch $c$ encountered conflict 3 thus leading to the creation of branch $e$, etc,... Branch $g$ encountered conflict 6 but the pair in conflict with $(16, 22)$ was encountered late in the exploration: the pair $(6, 2)$ has already been encountered and rejected. Thus the interdiction of forming $(6, 2)$ is transferred from branch $g$ to the new branch $h$ in addition of $(16, 22)$ which is directly involved in the conflict and $(14, 20)$ since the corresponding node in the EBT is higher than the one corresponding to $(16, 22)$.

**Algorithm 255:** `BuildExplorationBinaryTree`$(B, \alpha)$

**Data:** A set of branches $B$ and a starting point $\alpha$
**Result:** The Exploration Binary Tree $T$

```
T ← ∅
T.root ← α
for β ∈ B do
    cursor ← T.root
    went_left ← False
    for (pair, event) ∈ o_{α,β} | pair ∉ α ∧ event ∈ {A, F} do
        if went_left then
            if cursor.left == None then
                cursor.left ← (g, h)
        else
            if cursor.right == None then
                cursor.right ← (g, h)
        if event == A then
            went_left ← False
        else
            went_left ← True
    end
    if went_left then
        cursor.left ← b
    else
        cursor.right ← (g, h)
end
return T
```

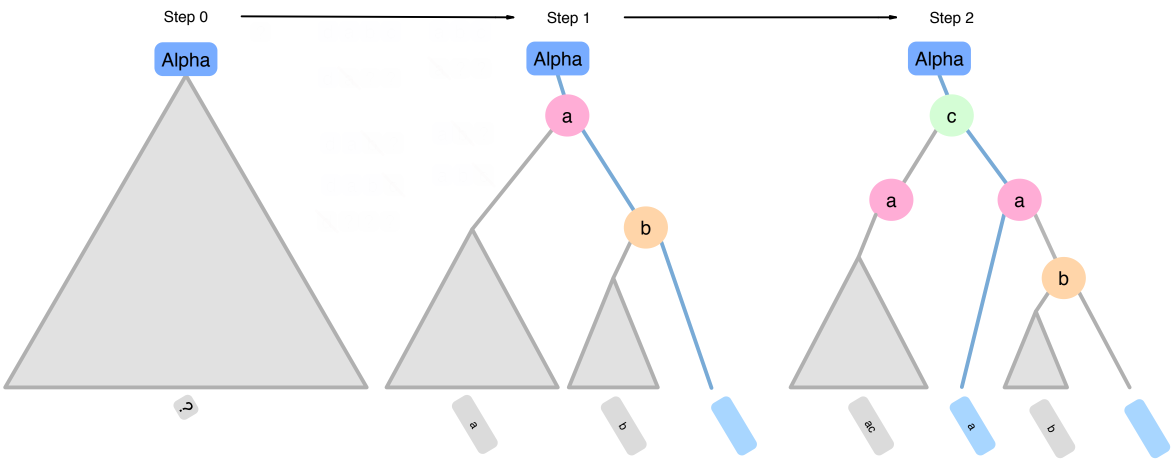

Figure 1.4: **Tree view of the branching-backtrack mechanism:** This display depicts the very same situation presented as an exemple in figure 1.1 and offers the opportunity to illustrate how the exploration of a new branch reveals a part of the EBT. At the beginning (Step 0), nothing is known about the EBT except for $\alpha$ being the root. After the exploration of the first branch $(\alpha, \beta = \epsilon)$ (Step 1, in light blue), two conflicts have been encountered involving pairs $a$ and $b$. As a consequence, the exploration not only revealed a path from the root to a leaf (in light blue) but also that the nodes corresponding to $a$ and $b$ have a left child in the EBT since the two new branches will pass through them. We then explore the branch $(\alpha, \beta = \{a\})$ (Step 2) and reveal another path from the root to a leaf (in light blue) and the node corresponding to pair $c$. The presence of a second node corresponding to pair $a$ is due to the fact that the new branch will also forbid $a$. This mechanism has been briefly explaining in figure 1.1 but is extensively covered in lemma 16(cf. second situation).

**Definition 5.** Two pairs $(g,h)$ and $(g',h')$ in $V_G \times V_H$ are called *exclusive* if there is no csg $S$ of $G$ and $H$ with the corresponding injections $i_G$ and $i_H$ such that there exists $x,y \in V_S$ such that

$$i_G(x) = g, i_G(y) = g', i_H(x) = h, i_H(y) = h'$$

Two exclusive pairs $(g,h)$ and $(g,h')$ (resp $(g,h)$ and $(g',h)$) are called *strongly exclusive.* Note that since $i_G$ and $i_H$ are injective, two pairs $(g,h)$ and $(g,h')$ (resp $(g,h)$ and $(g',h)$) in $V_G \times V_H$ are automatically exclusive.

For the remaining of this section, let $G$ and $H$ be two graphs.

**Theorem 6.** Given two maximal csg $S$ and $S'$ that share a common vertex of $G$ and $H$, with their corresponding inclusions $i_G, i'_G, i_H, i'_H$, then either $S$ and $S'$ are isomorphic or there exist $s \in V_S$ and $s' \in V_{S'}$ such that $(i_G(s), i_H(s))$ and $(i'_G(s'), i'_H(s'))$ are strongly exclusive.

*Proof.* Assume there is no such $s$ and $s'$. Define the following sets:

$$\begin{aligned}
V''_{com} &= \{(g,h) \mid \exists x \in V_S,\ \exists x' \in V_{S'}\ g = i_G(x) = i'_G(x') \text{ and } h = i_H(x) = i'_H(x')\} \\
V''_S &= \{(g,h) \mid \exists x \in V_S \text{ s.t. } g = i_G(x) \in i_G(V_S) \setminus i'_G(V_{S'}) \text{ and } h = i_H(x) \in i_H(V_S) \setminus i'_H(V_{S'})\} \\
V''_{S'} &= \{(g,h) \mid \exists x' \in V_{S'} \text{ s.t. } g = i'_G(x') \in i'_G(V_{S'}) \setminus i_G(V_S) \text{ and } h = i'_H(x') \in i'_H(V_{S'}) \setminus i_H(V_S)\} \\
V'' &= V''_{com} \cup V''_S \cup V''_{S'} \\
E''_l &= \{((g,h),(g',h')) \mid (g,g') \in E_{G,l} \text{ and } (h,h') \in E_{H,l}\} \\
E'' &= \bigcup_{l \in L} E''_l
\end{aligned}$$

**Lemma 7.** For all $x \in V_S$, there exists a unique $h \in V_H$ such that $(i_G(x), h) \in V''$.

*Proof.* To prove uniqueness, consider $h_1, h_2$ in $V_H$ such that $(i_G(x), h_1)$ and $(i_G(x), h_2)$ are in $V''$. By definition of $V''$, $h_j = i_H(x_j)$ $(j = 1,2)$ with $i_G(x_j) = i_G(x)$. Since $i_G$ is injective, we get $x_1 = x_2 = x$ and therefore $h_1 = h_2$.

To prove existence, let us show that $h = i_H(x)$ satisfies the condition. There are two cases to consider :

- Either $i_G(x) \notin i'_G(V_{S'})$, then $i_H(x) \notin i'_H(V_{S'})$. Indeed, if it was the case, this means that there is $x' \in V_{S'}$ such that $i_H(x) = i'_H(x')$. But then, the pairs $(i_G(x), i_H(x))$ and $(i'_G(x'), i_H(x))$ are strongly exclusive. But we assumed that there is no strongly exclusive pair.

- Or $i_G(x) \in i'_G(V_{S'})$, then consider $x' \in V_{S'}$ such that $i_G(x) = i'_G(x')$. Necessarily we have that $i_H(x) = i_H(x')$ (since otherwise the two pairs $(i_G(x), i_H(x))$ and $(i_G(x), i'_H(x'))$ would be strongly exclusive).

□

Note that we have found a candidate for such $h$ : $h = i_H(x)$.
There is a similar claim for $S'$.
Let $s \in V_S$ and $s' \in V_{S'}$ be the nodes that correspond to the shared vertex of $S$ and $S'$, i.e. $i_G(s) = i'_G(s')$ and $i_H(s) = i'_H(s')$.

**Lemma 8.** For every node $(g, h)$ in $V''$, there is an $E''$-path between $(g, h)$ and $(i_G(s), i_H(s))$.

*Proof.* Consider $(g, h)$ in $V''$. WLOG $g = i_G(x)$ and $h = i_H(x)$ for some $x \in V_S$. We know that there is a path in $S$ connecting $s$ and $x$. Let us note this path $s = x_0, x_1, ..., x_{n-1}, x_n = x$. By the first claim, for all $j$, $(i_G(x_j), i_H(x_j))$ is in $V''$. We know that both $i_G$ and $i_H$ respect edges and their labels. This means that for $\{x_j, x_{j+1}\} \in E_{S,l_j}$, $\{i_G(x_j), i_G(x_{j+1})\} \in E_{G,l_j}$ and $\{i_H(x_j), i_H(x_{j+1})\} \in E_{H,l_j}$.
By definition of $E''$, that means that for all $j$, $\{(i_G(x_j), i_H(x_j)), (i_G(x_{j+1}), i_H(x_{j+1}))\} \in E''$ which proves the claim. □

This proves that $(V'', E'')$ is a graph (it proves it is connected).

**Lemma 9.** The graph $(V'', E'')$ is a csg of $G$ and $H$.

*Proof.* We can define the following maps $i''_G : V'' \to V_G$ and $i''_H : V'' \to V_H$ as follows: $i''_G(g, h) = g$ and $i''_H(g, h) = h$.
These maps are indeed injective (by the uniqueness proved in the first claim).
Let us show that $i''_G$ respects edges ($i''_H$ is done in a similar fashion). Let $l$ be a label and let $((g, h), (g', h')) \in E''_l$. By definition of $E''_l$, $(g, g') \in E_{G,l}$, but we also have that

$(i''_G(g,h), i''_G(g',h')) = (g,g')$ which proves that $(V'', E'')$ is a subgraph of $G$ (and similarly of $H$). This means that $(V'', E'')$ is a csg of $G$ and $H$. □

**Lemma 10.** The graph $S$ is a subgraph of $(V'', E'')$.

*Proof.* This is done by the first claim. Define the map $i : V_S \to V''$ as $i(x) = (i_G(x), i_H(x))$. First this is an injection since $i_G$ is an injection.
Let us now consider an edge $\{x,y\}$ in $E_{S,l}$. Since both $i_G$ and $i_H$ respect edges and their labels, we have that $\{i_G(x), i_G(y)\} \in E_{G,l}$ and $\{i_H(x), i_H(y)\} \in E_{H,l}$. By definition, this means that $\{i(x), i(y)\} \in E''_l$.
However, remember that $S$ is a maximal csg of $G$ and $H$, which means that $S$ and $(V'', E'')$ are isomorphic. Since the proof is not specific to $S$ and can also be applied to $S'$, we have that $S'$ is also isomorphic to $(V'', E'')$ and therefore $S$ and $S'$ are isomorphic which proves the lemma. □

This concludes the proof of theorem **??**. □

This proof also justifies that we are going to use the following notation from now: a node $x$ of a subgraph of $G$ and $H$ will be denoted $(g,h)$ where $i_G(x) = g$ and $i_H(x) = h$.

**Definition 11. Paths in common subgraphs :** Given $x, y \in V_G \times V_H$, a *subgraph path from $x$ to $y$* denoted $x \to y$ is a finite sequence $(n_i)_{i \in \{0,\ldots,m\}}$ in $V_G \times V_H$ such that there exists a subgraph $S$ of graphs $G$ and $H$ such that $(n_i)$ is a path with $n_0 = x$ and $n_m = y$ that does not contain any loop in $S$.

We have that a finite sequence $((g_i, h_i))_{i \in \{0,\ldots,m\}}$ is a subgraph path $x \to y$ if, and only if,

1. for all $i \neq j$, $g_i \neq g_j$ and $h_i \neq h_j$,

2. for all $i \leq m-1$, there is a label $l$ such that $\{g_i, g_{i+1}\} \in E_{l,G}$ and $\{h_i, h_{i+1}\} \in E_{l,H}$,

3. $(g_0, h_0) = x$ and $(g_m, h_m) = y$

Indeed, note that such a subgraph path $(n_i)_{i \in \{0,\ldots,m\}}$ yields a csg $S$ of $G$ and $H$ where $V_S = \{n_i \mid 0 \leq i \leq m\}$ and $E_{l,S} = \{\{n_i, n_{i+1}\} \mid \{g_i, g_{i+1}\} \in E_{l,G} \text{ and } \{h_i, h_{i+1}\} \in E_{l,H}\}$ where $n_j = (g_j, h_j)$ for all $j$.

**Definition 12. Paths from $\alpha$ in common subgraphs :** Given $x \in V_G \times V_H$ and a starting point $\alpha = (n_a, n_b) \in (V_G \times V_H) \cup (V_G \times V_H)$, a *subgraph path from $\alpha$ to $x$* denoted $\alpha \rightarrow x$ is a subgraph path $(n_i)_{i \in \{0,\dots,m\}}$ such that $n_m = x$ and either $n_0 = n_a$ and $n_1 = n_b$, or $n_0 = n_b$ and $n_1 = n_a$.

Similarly to the case of a subgraph path $x \rightarrow y$, such a subgraph path $\alpha \rightarrow x$ also yields a csg containing the node $x$ and the a starting point $\alpha$.

**Definition 13. $p$Path :** A *ppath* $\alpha \rightharpoonup (g', h')$ is a path from $\alpha \rightarrow (g'', h'')$ in a csg $S$ such that the node $(g'', h'')$ is in the neighbourhood of node $(g', h')$ in the csg $S$.

An exploration that added the pairs in a $p$path $\alpha \rightharpoonup (g', h')$ to its associated mapping will necessarily consider $(g', h')$ and, if $(g', h') \notin \alpha$, add it if possible or otherwise raise a conflict $((g', h'), P)$ with $P$ the pairs already added and strongly exclusive with $(g', h')$. This notion enables us to describe pairs encountered during the exploration but not added to the mapping. Those pairs are crucial regarding conflict management as will be shown in theorem 14 and lemmas 15 and 16.

**Theorem 14.** For all maximal common subgraph $S$ of graphs $G$ and $H$, for all starting point $\alpha$ in $S$, $S$ is in the set of maximal common subgraphs found by the algorithm with ($G$,$H$,$\alpha$) as input.

*Informal intuition of the strategy* We will navigate in the EBT by bifurcating according to the content of $S$ (and proving that it is always possible to do so) until we reach a leaf. We then show that the mcsg corresponding to this leaf is isomorphic to $S$ using theorem 6.

*Proof.* Let us consider an arbitrary mcsg $S$ of $G$ and $H$. Let us also consider an arbitrary starting point $\alpha \in S$, let $M_\alpha$ be the set of mcsg of $G$ and $H$ constructed by the algorithm starting from $\alpha$.

We start at the root of the EBT which corresponds to $\alpha$ and will recursively go down in the tree using a pointer that we initiate as the right child of the root (the root only has a right child and never has a left child). We recall that $\rho$ denotes the sequence of decisions made from the root of the EBT.

Let us call $\rho(g,h)$ the node of the EBT the pointer is currently on. There are two possibilities.
1. Either $(g,h) \notin S$ and we move the pointer to the left child of $\rho(g,h)$. Since $(g,h) \notin S$, $S$ contains both a pair $(g',h')$ strongly exclusive with $(g,h)$ and a path $\alpha \to (g',h')$ and thus a $p$path $\alpha \rightharpoonup (g',h')$. Both those paths are compatible with context $\rho$ since $S$ is. As a consequence $\rho(g,h)$ has a left child by lemma 15.
2. Either $(g,h) \in S$ and we move the pointer to the right child of $\rho(g,h)$. Since $S$ is naturally compatible with the sequence of decisions $\rho$ and $(g,h) \in S$ thus $\rho(g,h)$ has a right child by lemma 16.
By repeating this process we ultimately reach a leaf of the EBT which corresponds to a common subgraph $S' \in M_\alpha$. By construction, $S'$ cannot be extended except maybe by adding a forbidden pair. However all pairs forbidden during the production of $S'$ are not in $S$. As a consequence the set of pairs $(g,h) \in S'$ is exactly the set of pairs $(g,h) \in S$ thus $S'$ and $S$ are the same mcs and $S$ is in $M_\alpha$ using theorem 6.

**Lemma 15.** For all nodes $\rho(g,h)$ in the EBT such that there exists a $p$path $\alpha \rightharpoonup (g',h')$ with $(g',h')$ strongly exclusive with $(g,h)$ and $\alpha \rightharpoonup (g',h')$ compatible with the decisions in $\rho$, then $\rho(g,h)$ has a left child.

*Proof.* Since a non-leaf node without a right child always has a left child, we will naturally assume that a node has a right child when trying to prove said node has a left child.
We will proceed by recursion on the number $n$ of pairs that need to be forbidden in order to form $\alpha \rightharpoonup (g',h')$.
If $\alpha \rightharpoonup (g',h')$ can be formed in the rightmost leaf from $\rho(g,h)$ (in other words $n = 0$), then the corresponding exploration has necessarily considered $(g',h')$ while $(g,h)$ was already in the mapping thus leading to a conflict and the creation of a left child to $\rho(g,h)$. This is our base case.
Let us now assume that the claim is true when $n$ pairs need to be forbidden and consider a situation in which the formation of $\alpha \rightharpoonup (g',h')$ requires $n+1$ pairs to be forbidden. A visual support for this induction step is provided in figure 1.5.
Let us consider $(g_n,h_n)$, the first of those pairs according to the exploration order

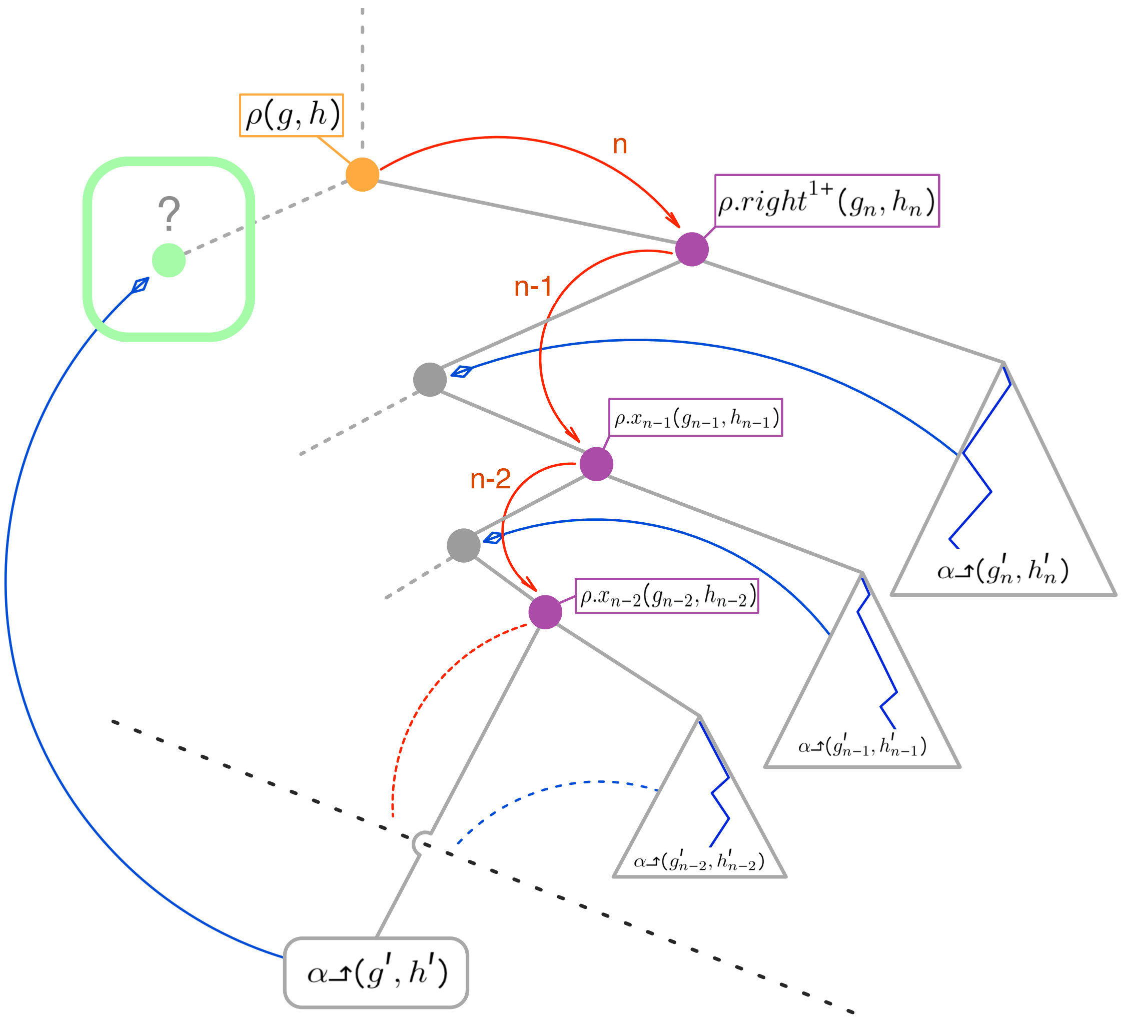

Figure 1.5: **Visual support for lemma 15:** This figure provides an informal visualization of the recursive pattern used in proof of lemma 15. Our goal is to prove the existence of a left child (in green in the figure) of node $\rho(g, h)$ (in orange). To do so we will descend in the subtree having the right child of $\rho(g, h)$ as its root, looking for $\alpha{\uparrow}(g', h')$ with $(g', h')$ a pair strongly exclusive with $(g, h)$. We know such $p$path exists thanks to $S$ but we still need to show it can be formed in this subtree as it is necessary for $\rho(g, h)$ to have a left child. The pairs that are exclusive with pairs in $\alpha{\uparrow}(g', h')$ are displayed in violet. Each such pair corresponds to a step in the descent represented in red. Those pairs need to be forbidden and thus the descent needs to pass through their left child (in grey). As a consequence, we apply the recursion to each of them, thus finding a suitable $p$path in the subtree having their right child as the root and thus guaranteeing the existence of their left child (guarantees represented by the blue arrows). Ultimately, we reach a leaf corresponding to a branch that forms $\alpha{\uparrow}(g', h')$ thus proving that $\rho(g, h)$ has a left child.

and $\rho.right^{1+}(g_n, h_n)$ the corresponding node in the EBT. Let us first assume that $\rho.right^{1+}(g_n, h_n)$ has a left child. Descending into its left child means forbidding $(g_n, h_n)$ thus leaving only $n$ pairs to be forbidden to form $\alpha \lrcorner(g', h')$. Let us now prove that $\rho.right^{1+}(g_n, h_n)$ has a left child. Since $\rho.right^{1+}(g_n, h_n)$ is the first pair strongly exclusive with a pair $(g'_n, h'_n)$ in $\alpha \lrcorner(g', h')$ encountered from $\rho(g, h)$, the $p$path $\alpha \lrcorner(g'_n, h'_n)$, prefix of $\alpha \lrcorner(g', h')$, is compatible with context $\rho.right^{2+}$ (i.e. it can be formed from the right child of $\rho.right^{1+}(g_n, h_n)$) and its formation requires $m$ pairs to be forbidden with $m \leq n$. By recursion $\rho.right^{1+}(g_n, h_n)$ has a left child.
We can then look for the next pair to be forbidden and repeat the processus up to the point where there are none remaining. The corresponding leaf will necessary form $\alpha \lrcorner(g', h')$. By doing so we will need several nodes to have a left child just like $\rho.right^{1+}(g_n, h_n)$ but the recursion will guarantee they do since the number of pairs to be forbidden to form the corresponding $p$path will decrease each time and always be smaller or equal to $n$.
As a consequence, any left child we might need to pass by during the recursive descent of the EBT is guaranteed to exist. □

**Lemma 16.** For all nodes $\rho(g, h)$ in the EBT such that there exists a maximal common subgraph $S$ compatible with decisions in $\rho$ and $(g, h) \in S$, then $\rho(g, h)$ has a right child.

*Proof.* Any eligible pair is added by the algorithm by default and thus granting a right child to the corresponding node in the EBT. However, there are two situations that may create a non-leaf node without a right child in the EBT.
The first situation occurs when encountering a conflict $((g, h), P)$ with $P = \{ (g', h'), (g'', h'') \}$. Let us assume without loss of generality that $(g', h') <_{o_{\alpha,\beta}} (g'', h'')$, in this case the EPT will contain two nodes $\rho(g', h')$ and $\rho.left.x(g'', h'')$ (with $x = right^*$) corresponding to the two pairs in $P$. The node $\rho.left.x(g'', h'')$ might not have a right child. However:

- either there are no mcs compatible with the sequence of decision $\rho.left.x.right$ and there is no need for $\rho.left.x(g'', h'')$ to have a right child.

- or there is at least one mcs $S'$ compatible with the sequence of decision $\rho.left.x.right$. The sequence of decision $\rho.left.x.right$ implies that $(g', h') \notin S'$ and $(g'', h'') \in S'$. Moreover $(g'', h'') \in S'$ implies that $(g, h) \notin S'$ as $(g'', h'')$ strongly exclusive with

$(g,h)$. Finally, $(g',h') \notin S'$ and implies that $S'$ contains a pair $(g^\dagger, h^\dagger)$ strongly exclusive with $(g',h')$ but not with $(g'',h'')$ since $(g'',h'') \in S'$. $S'$ is connected so it contains a path $\alpha \rightarrow (g^\dagger, h^\dagger)$ compatible with $\rho$ and so a $p$path $\alpha \rightharpoonup (g^\dagger, h^\dagger)$ compatible with $\rho$. Thus a branch in the right child of $\rho(g',h')$ will discover a conflict $((g^\dagger, h^\dagger), P)$ with $(g',h') \in P$ and $(g'',h'') \notin P$. Let us start with the simplest case: $P = \{(g',h')\}$. The branch created from this conflict will follow the sequence of decision $\rho.left.x.right$ thus leading to the creation of a right child to $\rho.left.x(g'',h'')$. Now let us consider $P = \{(g',h'),(g^\diamond,h^\diamond)\}$. Since $(g^\diamond,h^\diamond) \notin S'$ and $S'$ compatible with $\rho.left.x.right$, forbidding $(g^\diamond,h^\diamond)$ will not impact the sequence of decision and will also lead to the creation of the right child.

The second situation may happen during the creation of a branch. A visual support for this situation is provided in figure 1.6.

Let us consider a new branch $(\alpha, \beta')$ created from branch $(\alpha, \beta)$ to solve conflict $((g,h),P)$. Any pair $(g',h') \in \beta'$ inherited from $\beta$ which has been encountered between a pair in $P$ and $(g,h)$ (according to $o_{\alpha,\beta}$) may correspond to a non-leaf nodes without a right child in the EBT (as well as the second pair in $P$ has described in the firs situation). The second situation can be considered a generalization of the first one as it may produces several nodes without a right child but only if no mcs is compatible with a sequence of decisions that would lead to any of the missing right childs. This can be proven for all nodes created without a right child by repeating the proof of the first situation for all of those nodes starting by the one closest to the root. Let us denote $(g_1,h_1)$ the first pair in $P$ (according to $o_{\alpha,\beta}$) and use two pointers $a$ and $b$. We start with $a = \rho(g_1,h_1)$, the node of the EPT created to solve the conflict $((g,h),P)$ and $b = \rho.left.x(g_2,h_2)$ the first new node of the EPT without a right child. We then apply the proof of the first situation replacing $(g',h')$ with $a$ and $(g'',h'')$ with $b$. According to the proof, either $b$ should have a right child and will be created one or $b$ does not need one. We move pointer $b$ to the next node without a right child and if a right child is created for $b$ we move pointer $a$ so $a = b$. We then repeat this procedure until all nodes created without a right child have been covered.

As a consequence, and since $S$ is a mcs, any right child we might need to pass by during

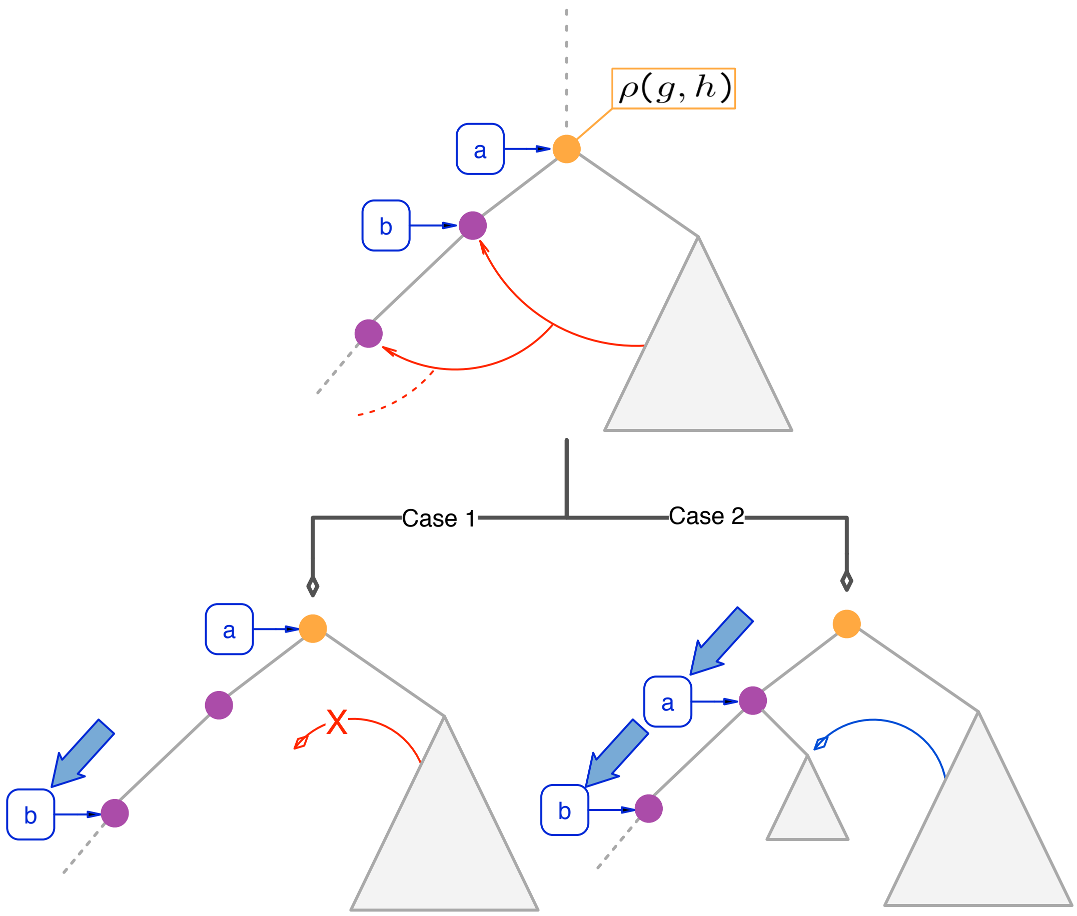

Figure 1.6: **Visual support for the second situation described in lemma 16:** This figure provides an informal visualization of the second situation described in lemma 16 and of the corresponding proof. The upper subtree displays the situation after the creation of the branch $(\alpha, \beta')$ because of a conflict encountered in the right subtree figured by a triangle. Several nodes in violet have been created without a right child. We initiate the pointer and move on one of the two cases figured below, depending on the outcome. If the first node without a right child does not need one, we go into case 1 and move the pointer $b$ to the next node without a right child. If the first node without a right child needs one, we go into case 2 and recursively prove that the right child will be created in another branch, and thus can move both pointers.

the recursive descent of the EBT is guaranteed to exist. □

As we are able for any maximal common subgraph to find a leaf in the EBT corresponding to it, all maximal common subgraph of $G$ and $H$ are found by the algorithm. □

##### 1.4.4 Efficiency

The complexity of the exploration of a single branch is in $\mathcal{O}(|L| \times n)$ with $n$ the size of the common subgraph produced. As $|L|$ the size of the set of colors of the edges is fixed, we consider an exploration to be linear in the size of the output and thus optimal.
Since $\bigcap_{\forall \alpha} mcsg_{G,H,\alpha} \neq \emptyset$, there is an overlap in the set of mcs produced. However, the cost of this overlap is bounded by a $n^2$ factor which may be considered negligible compared to the exponential size of the output.
Finally, and this is the greatest issue, the number of branches created to output a given $mcsg_{G,H,\alpha}$ may be greater that the number of mcsg in it. Indeed, in order to ensure correction, the algorithm creates all branches that may output a mcsg (as proven with theorem 14) but it also produces branches that end up producing a csg that is not maximal. This issue could happen in two situations described in the two next paragraphs.

**Pointless conflicts:** We call *pointless conflict* a conflicts $((g,h),P)$ such that there is no valid path $\alpha \rightarrow (g,h)$. To put it differently, the pair $(g,h)$ is reached by the algorithm thus there is at least one $p$path $\alpha \Lsh (g,h)$ but all the $p$path contain a pair in $P$. As a consequence, forbidding the pair in $P$ in order to allow the formation of $(g,h)$ also cuts all paths to reach $(g,h)$ and so is pointless.
A branch created from a pointless conflict is very likely to produce a common subgraph that is not maximal (in general, a subgraph of the one produced by its mother branch). However, checking if given conflict is pointless or not cannot be done faster than by exploring the branch created from this conflict.

**Inheritance of forbidden pair:** When a new branch $\{\alpha, \beta'\}$ is created from a branch $\{\alpha, \beta\}$ because of conflict $((g,h),P)$, it may inherit pairs from $\beta$ encountered after the pairs in $P$ but before $(g,h)$ as mentioned in section 1.4.2, figure 1.1 and figure 1.4 as well as in lemma 16 (second situation).
We indeed assume that the interdictions enforced before encountering $(g,h)$ may be required to ever reach $(g,h)$. However, if such an interdiction is not required, the interdiction may be either pointless (the forbidden pair may be never encountered and thus the corresponding interdiction has no impact on the exploration) or even detrimental,

causing the branch to produce a common subgraph which is not maximal.
However, this mechanism produces all the branches needed to output all mcsg (cf. lemma 16) while offering a crucial property: branches that produces non-maximal common subgraphs never creates new branches. This property of the algorithm drastically limits the number of non-maximal branches generated.

**Test of maximality:** Testing if a common subgraph $S$ outputted by branch $\{\alpha, \beta\}$ is maximal can easily be done by checking for all pairs $(g, h) \in \beta$ encountered (yet rejected) during the exploration, if the common subgraph contains a pair strongly exclusive with $(g, h)$ (using theorem 6).

- If it does not, that implies that a strictly bigger common subgraph $S'$ can be produced by adding the pair $(g, h)$ to $S$. $S'$ is connected since $(g, h)$ has been encountered during the exploration. Thus $S$ is not maximal.
- If it does for all pairs $(g, h) \in \beta$, $S$ is maximal. Indeed, the exploration is only blocked by the interdictions contained in $\beta$. However, all pairs in $\beta$ are strongly exclusive with at least a pair in $S$ thus is maximal.

**Complexity:** To provide a meaningful complexity for this algorithm, we would need a way to count or to bound the number of occurrences of the two events we just mentioned. More generally it would require to express the number of conflicts in relation to the length of either the input or the output. We failed to find such formula despite our best efforts. It seems to be considerable work and out of the scope of this thesis.
Comparative unbiased benchmarks are also not an option because of the lack of both a balanced generic benchmark and similar algorithms to be compared to.
However, an application of this algorithm is presented in the next chapter in a specific context. We used this opportunity to provide some assessment of its performance. This solution is far from being satisfactory and we are currently looking for an alternative.
Without going into details to avoid redundancy with chapter 2, let us just mention that the performances in this context are very satisfying.

#### 1.5 Adaptation to other graphs

##### 1.5.1 Directed graphs

The algorithms have been described using undirected graphs for the sake of readability as it greatly simplifies many notations. However, the algorithms can easily be extended to directed edge-labelled graphs respecting some properties.

We first have to adapt the definition of *proper edge coloring* to directed graphs. We thus will use the classical definition of *proper edge coloring* and consider that the colors on the edge of graph $G$ form *proper edge coloring* if all edges incident to a same node have different colors i.e. $\forall g, g^{'}, g^{''} \in V_G$ such that $\{g^{'}, g\} \in E_{l,g} and \{g^{''}, g\} \in E_{l^{'},g}$ we have $l \neq l^{'}$.

We then need to consider connectivity. Our strategy requires that all pairs $(g, h)$ can be accessed from any starting point $\alpha$ and thus that for all edge $\{g_1, g_2\}$ there exists an edge $\{g_2, g_1\}$. Moreover it requires that the if an edge $\{(g_1, h_1), (g_2, h_2)\}$ is in a common graph $S$, the edge $\{(g_2, h_2), (g_1, h_1)\}$ is also in $S$.

We haven't mentioned the coloring of the edges in the previous paragraph. We do not require that the reversed arrow has the same label. However, we do require a weaker condition: there exists a bijection $b : L \to L$ such that $\forall \{g, g^{'}\} \in E_{l,G}$, $\exists \{g^{'}, g\} \in E_{b(l),G}$. Please also note that working on such directed graphs is advantageous for us as the directions (if $l \neq b(l)$) provide additional constraints that limits the number of possibilities to explore.

##### 1.5.2 Extension to other graphs

The following graphs fall out of the scope of this work. As a consequence, we will only mention ideas to accommodate such graphs.

###### Other directed graphs

Some directed graphs $G = \{V, E, L\}$ that does not respect the three properties mentionned earlier may still be handled by manipulating the labels to obtain a different graph $G^{'} = \{V, E^{'}, L^{'}\}$ that respects the properties.

For instance, a graph $G$ that contains one or several $\{g_1, g_2\} \in E_{l,G}$ but no edge $\{g_2, g_1\}$

can be transformed into another $G'$ by adding a new label $l' \notin L$ and edges $\{g_2, g_1\} \in E_{l',G}$. Similarly, it might be possible to create a graph respecting the properties from a graph without a bijection $b : L \to L$ such that$\forall \{g, g'\} \in E_{l,G},\ \exists \{g', g\} \in E_{b(l),G}$ by creating new labels (for instance labels $l \wedge l'$ or $l \vee l'$) and/or new edges.

##### Multigraphs

Our algorithms is not designed to handle multigraphs, however some situations may still be managable.
Let us consider two connected and undirected edge-labelled multigraphs $G = \{V_G, E_G\}$ and $H = \{V_H, E_H\}$. Let's assume $\exists g_1, g_2 \in V_G \mid \exists \{g_1, g_2\} \in E_{G,l_a} \wedge \exists \{g_1, g_2\} \in E_{G,l_b}$.

If $g_1, g_2$ can be matched to any $h_1, h_2 \in V_H \mid \exists \{h_1, h_2\} \in E_{H,l_a} \vee \exists \{h_1, h_2\} \in E_{H,l_b}$, a solution is to replace all edges $\left(\{x_1, x_2\} \in E_{X,l_a} \cup E_{X,l_b}\right)$ with a new edge $\{x_1, x_2\}$ labelled $l_{a \vee b}$.

If $g_1, g_2$ can be matched to any $h_1, h_2 \in V_H \mid \exists \{h_1, h_2\} \in E_{H,l_a} \wedge \exists \{h_1, h_2\} \in E_{H,l_b}$, a solution is to replace all pairs of edges $\{x_1, x_2\} \in E_{X,l_a}, \{x_1, x_2\} \in E_{X,l_b}$ with a new edge $\{x_1, x_2\}$ labelled $l_{a \wedge b}$.

Those operations are to be repeated until the creation of two graphs $G'$ and $H'$ which are not multigraphs and thus can be handled by the algorithms provided they respects the properties mentioned previously.
This notion is obviously to be extended and adapted according to the specificities of the graphs and of the problem.

##### Node-labelled graphs

If the graphs are both node-labelled and edge-labelled, the labels on the node are just an extra condition to enforce :
$g \in V_{G,l_g}$ can be matched to $h \in V_{H,l_h}$ only if $l_g = l_h$, with $V_{X,l}$ the subset of nodes of graph $X$ which label is $l$.

### Chapter 2

### Extraction of Recurrent Structural Elements

#### 2.1 Management of exceptions to the proper edge-coloring

The annotation method of RNA structures obtained from biological experiments sometimes produces nodes involved in two base pairs of the same type. The labels of the edges of RNA 2D structure graphs containing such nodes do not form a *proper edge colouring* which is problematic since we want to apply the algorithms presented in chapter 1. Those violations might be artifacts from the interaction prediction method which is based on distance thresholds between the nucleobases' atoms. However, it is possible that those double interactions are actually biologically relevant and we thus propose a solution to handle them. From our observations, such node are very uncommon (a dozen nodes over the fifty-two thousands nodes of the complexes from the non-redundant RNA database maintained on RNA3DHub [5]). Moreover, in all the cases we observed, the nucleobases forming the same interaction twice was forming it with two other nucleobases consecutive in the backbone. As a consequence, we handle those exceptions by duplicating any connected component containing a problematic node as follows :

With $a, b, c \in N$ s.t. $\{a, b\}, \{a, c\} \in E_l$ and $b < c$, and $\kappa$ the connected component containing them. We create $\kappa_a^!, \kappa_a^-, \kappa_a^+$ and $\kappa_a^*$ with :

- $\kappa_a^! = \kappa - \{\{a, b\}, \{a, c\}\}$
- $\kappa_a^- = \kappa - \{\{a, c\}\}$
- $\kappa_a^+ = \kappa - \{\{a, b\}\}$
- $\kappa_a^* = \kappa \; with \; \{a, b\} \in E_{l^-}, \{a, c\} \in E_{l^+}$

Please note that $\{a, b\}$ and $\{a, c\}$ can always be compared by comparing the position of $b$ and $c$ in the backbone. $\kappa_a^-$ is thus the version of $\kappa$ where the "highest" edge of the two has been removed while $\kappa_a^+$ is thus the version of $\kappa$ where the "lowest" edge of the two has been removed.
The labels of the edges of those four versions of $\kappa$ form a *proper edge colouring* and represent three different interpretations of those exceptions :

- $\kappa_a^!$ : both interactions are artifacts
- $\kappa_a^-, \kappa_a^+$ : one interaction is valid, the other is an artifact
- $\kappa_a^*$ : both interaction are valid, as a consequence those triangular structures are meaningful and thus should only be matched to similar structures

Those interpretations will impact how we handle several violations occurring in the same connected component. Let $a$ and $x$ two nodes that are incompatible with a *proper edge colouring* in the same connected component $\kappa$. $\kappa^!$ and $\kappa^*$ will just stack leading to $\kappa_{ax}^!$ and $\kappa_{ax}^*$. However $\kappa^-$ and $\kappa^+$ need to cover every possibility leading to $\kappa_{ax}^{--}, \kappa_{ax}^{-+}, \kappa_{ax}^{+-}$ and $\kappa_{ax}^{++}$. This system is obviously extendable to any number of violation. Most observed violations are in separate connected components with only three of them being in the same.
This solution of duplicating some connected components does not rule out any possibility and thus offers the opportunity to compare the structures observed in each version. Moreover, the limited number of problematic nucleobases observed makes the cost negligible.

To avoid duplicating results, if the same structure is found in several versions of $c$, we only keep one occurrence. The conserved occurence is picked according to the following order: $\kappa^! > \kappa^- > \kappa^+ > \kappa^*$. This order prioritizes the connected component our solution impacted the least so the fact that an occurence is mentioned has found in $\kappa^*$ implies that the triangular structure was required. We also do not search common structure between different versions of the same component as they represent the same nucleotides of the same RNA.
Please note that we have been considering connected component in this section because $f'$ may disconnect the RNA 2D structure graphs.

#### 2.2 Gathering of partial results

The core of the process of transforming a set of maximal common subgraphs (mcsg) into a collection of recurrent interaction networks (RINs) relies on the application of the filtering function $f_{RIN}$ to each mcsg in order to obtain the RINs inside it. We thus obtain a set of RINs from each mcsg found. We merge those sets of RINs so identical RINs (i.e. which canonical graphs are isomorphic) are merged into a single RIN combining all their occurrences (without duplicates). However, the *set of sets RINs found in the mcsgs* we obtain is not identical to the *set of RINs found in the dataset* we are seeking. To obtain the later, we have to correct two issues in the former.

A. First, the *set of sets RINs found in the mcsgs* may contain RINs with incomplete collections of occurrences. To put it differently, those RINs can be found in the dataset at positions that are not covered by any occurrences in their respective collections of occurrences. Those missing positions have been captured by the mcsgs but were "consumed" by/for another RIN. For instance, let us consider two graphs $G$ and $H$ and two RINs $a$ and $b$ such that:
$1_A$. the canonical graph of $a$ is a subgraph of the canonical graph of $b$
$2_A$. $b$ has two occurrences in $H$ and one in $G$ while $a$ has two in both
Please note that because of $1_A$, each occurrence of $b$ induces an occurrence of $a$. Let us consider the occurrence of $a$ in $G$ that is not induced by an occurrence of $b$. This

occurrence will be captured by at least two mcsgs, one for each of the occurrences of $a$ in $H$. However the second occurrence of $a$ in $G$ may only be captured by mcsgs that are capturing the instances of $b$ at the same time. As a consequence, the output of $f_{RIN}$ when we applied it to those mcsgs will only contain $b$ and not $a$. As a result, $a$ might appear with only three occurrences instead of four.

However, this issue can easily be covered by checking the collections of occurrences of any such pairs of RINs $a$ and $b$ and creating any missing occurrence of $a$ from the occurrences of $b$.

B. Second, the *set of sets RINs found in the mcsgs* may contain pairs of RINs $a$ and $b$ such that:

$1_B$. the canonical graph of $a$ is a subgraph of the canonical graph of $b$

$2_B$. each occurrence of $a$ is induced by an occurrence of $b$ (to put it differently, $a$ can only be found in the dataset inside occurrences of $b$)

In such case, we consider that $a$ does not provide any additional information to the collection and thus that it needs to be removed for the sake of the readability of the collection of RINs.

The presence of such RINs in the *set of RINs found in the mcsgs* is due to the counterintuitive fact that a *maximal* common subgraph may contain a *"non-maximal"* RIN i.e. a RIN which canonical graph is a subgraph of another RIN that can be found at the same positions in the two graphs (this other RIN is actually found at those very positions but by another mcsg). This happens when the nodes of the pairs of nodes needed to capture the larger RIN have been used to form other pairs of nodes: the resulting common subgraph is indeed maximal but those other pairs of nodes happen to have been eliminated by $f_{RIN}$, thus losing maximality.

As a consequence, we need to filter the *set of RINs found in the mcsgs* to eliminate such RINs. This could be done by checking the dataset for pairs of RINs $a$ and $b$ that satisfy both conditions $1_B$ and $2_B$ and remove $a$ but the fact that we completed the collections of occurrences to cover for issue A allows for a simpler solution. Indeed, once the collections of occurrences are completed, for all pairs of RINs $a$ and $b$ that satisfy condition $1_B$, the number of occurrences of $a$ cannot be less than the number of occurrences of $b$. Moreover, the number of occurrences of $a$ first will be greater than the number of occurrences of $b$, except if the pair also satisfies condition $2_B$. As a consequence, we can simply go through all pairs that satisfy $1_B$, compare their number of occurrences and eliminate $a$ if they are equal.

#### 2.3 Parallel computing

The pipeline has been designed to support parallel computing as the problem is greatly compatible with it. Indeed, the production of each set of maximal common subgraphs of two RNA 2D structure graphs is independent and can be processed separately. Thus we divide the work between $n$ cores between a *master* process et $n-1$ *workers*. The *workers* are provided with pairs of RNA 2D structure graphs by the *master* and send back the corresponding set of proto-RINs (i.e. they produce the corresponding set of maximal common subgraphs and then process it into the corresponding set of proto-RINs before sending it back). The *master* ensures that all pairs of RNA 2D structure graphs are processed and gather the results. Gathering the results includes the merging of partial results into the final collection. Rather than doing the merging in one go at the end of the computation which would let it with little to do when the *workers* are working, the *master* rather performs this task progressively as the *workers* send back their results.
All running times (i.e. original `CaRNAval`, $RIN^{ab}$ and $RIN^{a}$) are measured on the same machine (Intel(R) Xeon(R) CPU E5-2667 0 @ 2.90GHz, Ubuntu 16.0.4 with 23 cores, total physical memory of 792 gigabytes). The numbers we provide are total consumptions (i.e. the sums of CPU time consumed for all the core).

### Bibliography

[1] Scott Fortin. The graph isomorphism problem. Technical report, Citeseer, 1996.

[2] Jacobo Torán. On the hardness of graph isomorphism. *SIAM Journal on Computing*, 33(5):1093–1108, 2004.

[3] Brendan D McKay and Adolfo Piperno. Practical graph isomorphism, II. *Journal of Symbolic Computation*, 60:94–112, 2014.

[4] Eugene M. Luks. Isomorphism of graphs of bounded valence can be tested in polynomial time. *Journal of Computer and System Sciences*, 25(1):42 – 65, 1982.

[5] Anton Petrov. *RNA 3D Motifs: Identification, Clustering, and Analysis*. PhD thesis, Bowling Green State University, 2012.
